# Type I interferon primes the alveolar epithelium to receive reparative signals from tissue-resident macrophages

**DOI:** 10.64898/2026.06.10.731366

**Authors:** Alan Y. Baez Vazquez, Daisy A. Hoagland, Alexander O. Mann, Yunkang Lin, Susanna M. Dang, Max Hauptschein, Louison Thorens, Shahinoor Begum, Martha A. Castro, Patricia Rodríguez-Morales, Alicia Lai, Irving Barrera, Dawei Sun, Andrea Shehaj, Fei Chen, Christophe Benoist, Carla F. Kim, Ruth A. Franklin

## Abstract

Lung repair in response to viral infection requires integrated communication between epithelial and immune compartments, yet the impact of antiviral mediators on epithelial regenerative capacity remains poorly defined. Here, we demonstrate that type I interferon (IFN-I) signaling primes the lung for alveolar renewal following viral challenge. IFN-I contributes to the induction of an interferon-stimulated gene-high (ISG)^hi^ Sca-1^Pos^ population of alveolar type II (ATII) epithelial cells. Sca-1^Pos^ ATIIs exhibit enhanced proliferative capacity and increased organoid-forming efficiency compared with their Sca-1^Neg^ counterparts. Viral challenge concurrently drives phenotypic reprogramming of tissue-resident alveolar macrophages (trAMs). Sca-1^Pos^ ATIIs display heightened responsiveness to oncostatin M (OSM) and, following viral challenge, require trAM-derived OSM for their proliferation. Together, these findings reveal that viral stimuli induce coordinated IFN-I-dependent epithelial and macrophage states that poise the lung for regeneration, positioning IFN-I not only as a central antiviral defense mechanism but as a priming signal that prepares lung tissue for renewal.

## Introduction

The lung is continuously exposed to the external environment and is therefore particularly vulnerable to infection by pathogens. Viral respiratory infection elicits rapid activation of the immune system to eliminate the virus, resulting in tissue damage that can impair lung function. Successful recovery from infection requires precisely coordinated interactions between epithelial and immune compartments to repair injured tissue and restore homeostasis across a single epithelial layer that separates the host from the environment (*1*). Within the lung alveoli, the site of gas exchange, alveolar type II (ATII) cells are key drivers of epithelial regeneration through both self-renewal and differentiation into alveolar type I (ATI) cells (*2*). In settings of severe injury, airway progenitor populations, including club cells, basal cells, and bronchoalveolar stem cells (BASCs), can also contribute to alveolar repair (*3–7*). However, the early events that lead to alveolar proliferation and reconstitution during viral infection are not fully understood.

Previous work has highlighted the crucial role of macrophages in tissue repair across organs, including the lung (*8*). Several studies have shown that infiltrating monocytes and monocyte-derived macrophages play an important role in airway and alveolar repair, however the contribution of tissue-resident macrophages is less clear (*9–15*). With the increasing use of single-cell RNA sequencing (scRNA-seq) technologies to define macrophage heterogeneity after inflammatory challenges, monocyte origins of macrophage subsets are often inferred without direct validation using lineage tracing models. As a result, the extent to which phenotypic remodeling of tissue-resident macrophages contributes to epithelial repair remains unclear. Dissection of macrophage plasticity is essential for designing strategies to selectively reprogram or expand specific macrophage populations for therapeutic benefit in a variety of disease contexts.

Type I interferons (IFN-I) are central mediators of antiviral defense in the lung, rapidly produced by epithelial cells and resident macrophages upon infection to generate a protective antiviral state (*16–18*). In addition to restricting viral replication, IFN-I signaling modulates a variety of innate and adaptive immune functions by priming immune cells for more robust activation, cytokine production, and cytotoxicity, which are required to establish protective immunity (*18, 19*). Although IFN-I-mediated priming is well characterized in immune cells (*20*), the functional consequences of IFN-I exposure for epithelial cells within damaged tissues are not fully defined. In epithelial cell populations, IFN-I has been shown to promote cell death and suppress proliferation, leading to the prevailing view that IFN responses temporally precede, and may antagonize, tissue repair (*21–24*). However, emerging evidence suggests that repair programs are initiated rapidly and proceed in parallel with antiviral responses during infection (*11*). In this setting, where epithelial cells are transiently exposed to elevated IFN-I levels, we asked whether IFNI directly influences the regenerative capacity of alveolar epithelial cells during lung repair.

Using two *in vivo* models of acute IFN-I induction, sublethal influenza A virus (IAV) infection and local delivery of the viral mimic poly(I:C), we found that IFN-I signaling in the lung transiently induced a Sca-1^Pos^ ATII population that comprised the primary proliferative subset and facilitated epithelial renewal. Alongside these changes, IFNI exposure also induced phenotypic alterations in tissue-resident alveolar macrophages (AMs), representing profiles typically attributed to monocyte-derived cells. Functionally, Sca-1^Pos^ ATIIs exhibited enhanced sensitivity to macrophage-derived growth factors, enabling a heightened response to proliferative cues. In addition, Sca1^Pos^ ATIIs demonstrated increased organoid-forming efficiency *ex vivo* compared with Sca-1^Neg^ ATIIs, indicative of elevated progenitor potential and regenerative capacity. Together, these findings reveal a previously unrecognized role for IFN-I in coordinating epithelial-macrophage crosstalk to prepare the lung for renewal following inflammation and injury.

## Results

### Viral stimuli induce a proliferative ISG^hi^ Sca-1^Pos^ ATII cell state

To interrogate alveolar repair dynamics following viral infection, we performed scRNA-seq on epithelial and stromal lung fractions at 0, 7, 12, and 28 days post-infection (dpi) with sublethal IAV (A/PR/8/34) (Fig. 1A, fig. S1A and S1B, and table S1). Within ATIIs at baseline (Fig. 1B and 1C, fig. S1C, and table S2), we identified a major steady-state ATII population and a small subset of cells resembling alveolar differentiation intermediates (ADIs) (*25, 26*) (fig. S1D and table S3). At 7 dpi, we found ATIIs to be enriched for interferon-stimulated genes (ISGs), forming ISG^mid^ and ISG^hi^ clusters (Fig. 1C and 1D, and table S3). ISG^hi^ ATIIs were enriched for *Ly6a* (Stem cell antigen1, Sca-1) (Fig. 1D), both an ISG (*27, 28*) and canonical stem cell marker (*6, 28*). At 12 dpi, we observed a population of proliferating ATIIs and a population that was transcriptionally similar to previously described *activated* (*29*) or *primed* (*25*) ATIIs (Fig. 1C, fig. S1E, and table S3). Notably, the ISG^hi^ ATII subcluster detected at 7 dpi temporally preceded the appearance of activated ATIIs and exhibited an overlapping gene signature, suggesting a potential relationship between these states (fig. S1C and S1E). At 28 dpi, we observed the same populations present at baseline, but with an increased proportion of ADIs (Fig. 1C), consistent with their previously reported accumulation after lung injury (*22, 25, 26, 29, 30*).

**Figure 1.**
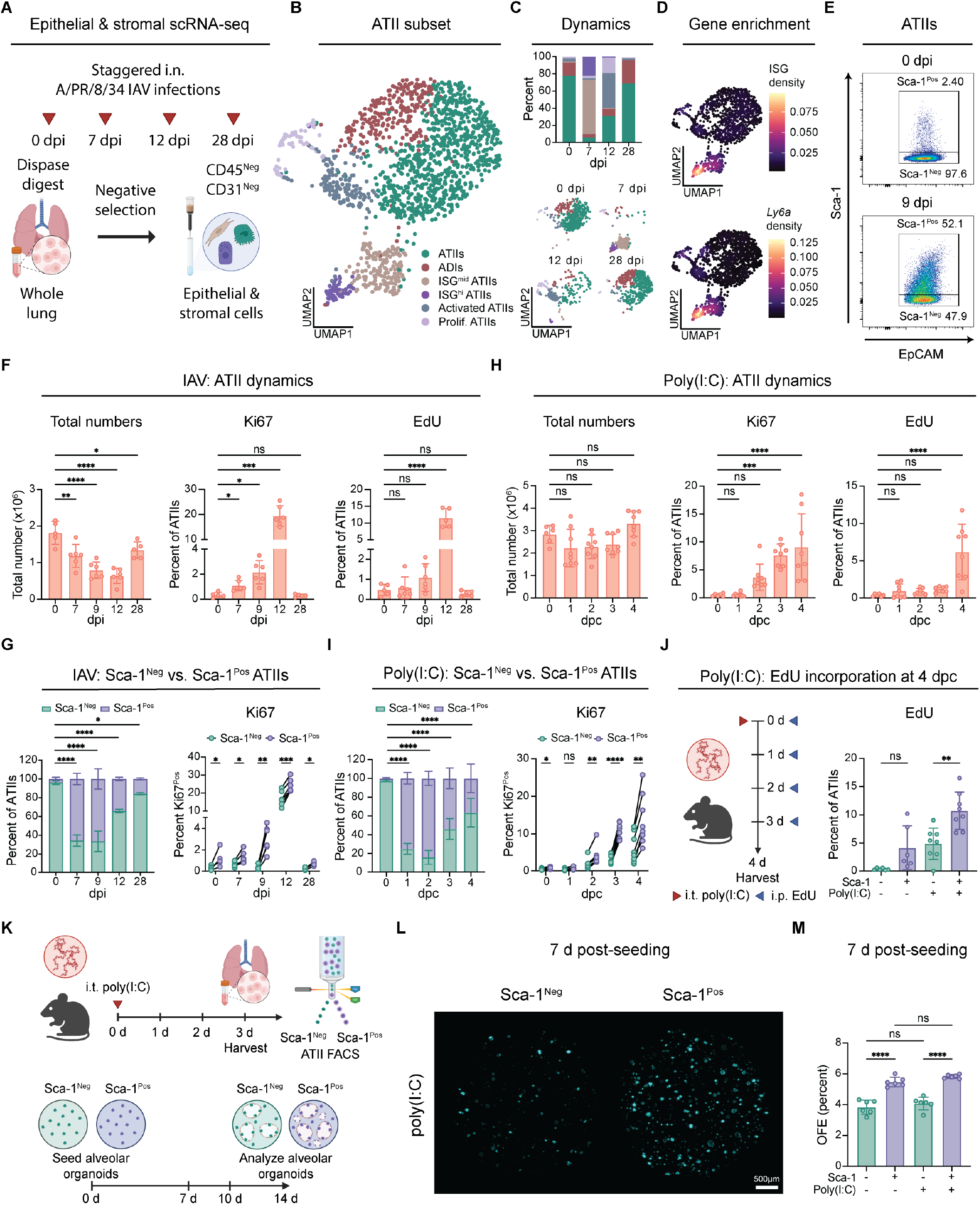
Exposure to viral stimuli induces a proliferative ISG^hi^ Sca-1^Pos^ alveolar epithelial cell state. **(A–D)** Single-cell RNA-seq was performed on lung epithelial and stromal cells following sublethal i.n. IAV infection (A/PR/8/34; 6.5 TCID_50_ for females and 32 TCID_50_ for males). Mice were infected and harvested at 0 (mock-infected), 7, 12, or 28 days post-infection (dpi; A; n = 1 male and 1 female mouse per group). Uniform manifold approximation and projection (UMAP) of ATII subset (B; 1,529 cells). Dynamics of ATIIs over time (C). ISG module enrichment and *Ly6a* gene expression nebulosa density plots (D). **(E)** Flow cytometry quantification of ATII total numbers or proliferation (Ki67 or EdU) after mock (0 dpi) or IAV infection (5 TCID_50_ ; n = 5 -6 mice per group; 7, 9, 12, or 28 dpi). Mice were injected i.p. with EdU (200 μg) 1 day before harvest. **(F)** Representative gating of Sca-1 expression on ATIIs at 0 or 9 dpi (5 TCID_50_ IAV) . **(G)** Flow cytometry quantification of Sca-1^Neg^/Sca-1^Pos^ ATIIs of total ATIIs and percent Ki67^Pos^ of Sca-1^Neg/Pos^ ATIIs. **(H)** Flow cytometry quantification of ATII total numbers or proliferation after mock (0 dps) or poly(I:C) (i.t. 33.75 μg) challenge (1, 2, 3 and 4 days post-challenge; dpc; n = 6 -8 mice per group). **(I)** Flow cytometry quantification of Sca-1^Neg^/Sca-1^Pos^ ATIIs of total ATIIs and percent Ki67^Pos^ of Sca-1^Neg/Pos^ ATIIs. **(J)** Percent cumulative EdU (200 μg per treatment) incorporation of Sca-1^Neg^/Sca-1^Pos^ ATIIs at 4 dpc. **(K–M)** FACS isolation and organoid characterization of Sca-1^Neg/Pos^ ATIIs from mock-or poly(I:C)-challenged (33.75 μg) mice (2 biological replicates and 3 technical replicates per group; quantified at 7, 10, and 14 days post-seeding; dps). Experimental scheme (K). Representative organoid image at 7 dps (L). Organoid forming efficiency (OFE) quantification at 7 dps (M). Data are representative of at least two independent experiments for (F, G, and K-L). Data from two independent experiments were pooled for (H-J). One-way ANOVA with Dunnet’s post-test for (F-J) and (M). Calculated on Sca-1^Pos^ ATIIs for fraction of total plots (G) and (I). Error bars indicate standard deviation (SD). \**p* ≤ 0.05, * * *p* ≤ 0.005, * * \**p* ≤ 0.0005, * * * * *p* ≤ 0.0001.

To confirm infection-induced ATII dynamics, we performed flow cytometry analysis across various timepoints following IAV infection (fig. S1F). The specificity of our gating strategy for ATIIs was confirmed by high pro-SFTPC staining and low CCSP (CC10), CD24, and EpCAM expression (fig. S1G). Following infection, ATII loss was evident at 7 dpi, decreased through 12 dpi, and largely resolved by 28 dpi (Fig. 1E). ATII proliferation was observed at 7 dpi and peaked at 12 dpi by both Ki67 positivity (Fig. 1E) and EdU incorporation (Fig. 1E and fig. S1H). We further confirmed that the Sca-1^Pos^ ATII population increased in magnitude by 7 dpi, appeared to peak at 9 dpi, and was largely resolved by 28 dpi (Fig. 1F and 1G). Despite the known anti-proliferative effects of IFN (*21–24, 31*), we observed that Sca-1^Pos^ ATIIs exhibited greater proliferative capacity than their Sca-1^Neg^ counterparts across all timepoints, as measured by Ki67 positivity (Fig. 1G).

Our scRNA-seq time course analysis implicated IFN signaling as a likely inducer of Sca-1^Pos^ ATIIs. To investigate the role of IFN-I in epithelial proliferation, we employed a single-dose acute poly(I:C) challenge model to induce IFN-I and mitigate potential artifactual enrichment of Sca-1^Pos^ ATIIs resulting from ATII depletion observed during IAV infection. One intratracheal (i.t.) treatment of poly(I:C) was sufficient to induce proliferation without significant ATII cell loss (Fig. 1H). As expected, poly(I:C) challenge was also sufficient to induce the Sca-1^Pos^ state within ATIIs (Fig. 1I). Like in IAV infection, Sca-1^Pos^ ATIIs were more proliferative than their Sca-1^Neg^ counterparts by Ki67 positivity (Fig. 1I) and by cumulative EdU incorporation at 4 dpc (Fig. 1J).

Given our findings in both IAV infection and poly(I:C) challenge models that Sca-1^Pos^ ATIIs exhibited greater proliferative capacity than Sca-1^Neg^ ATIIs, we next asked whether this population also displayed enhanced organoid-forming efficiency (OFE) *ex vivo*. We sorted Sca-1^Neg^ and Sca-1^Pos^ ATIIs from PBS-or poly(I:C)-challenged mice (fig. S2A) and generated alveolar organoid cultures (*32*), imaging the organoids at 7, 10, and 14 days postseeding (Fig. 1K and 1L, and fig. S2B). Organoids generated from Sca-1^Pos^ ATIIs showed enhanced OFE when compared to their Sca-1^Neg^ counterparts, independent of prior challenge (Fig. 1M, and fig. S2C). These data suggest that IFN-I exposure may augment the proliferative capacity of ATIIs.

### Viral challenge induces a transient CD11b^Pos^ CD11c^Pos^ AM state

To examine immune cell dynamics following sublethal IAV challenge, we performed cellular indexing of transcriptomes and epitopes by sequencing (CITE-seq) on immune cell fractions at 0, 7, 12, and 28 dpi (Fig. 2A, fig. S3A and S3B, and table S1). Within macrophages and monocytes (Fig. 2B, fig S3C, and table S2), we observed three major populations at baseline: AMs, interstitial macrophages (IMs), and monocytes (Fig. 2C). At 7 dpi, few AMs were observed, while a large transient population expressing both canonical AM and IM/monocytederived macrophage (moMac) markers emerged (CD11b^hi^ CD11c^lo^ macrophages) (Fig. 2C and 2D). At 12 dpi, we identified AMs, IMs, monocytes, putative monocytederived AMs (moAMs), and proliferating AMs (Prolif. AMs) (Fig. 2C). At 28 dpi, we observed a large fraction of AMs and a smaller fraction of monocytes, moAMs, and IMs (Fig. 2C).

**Figure 2.**
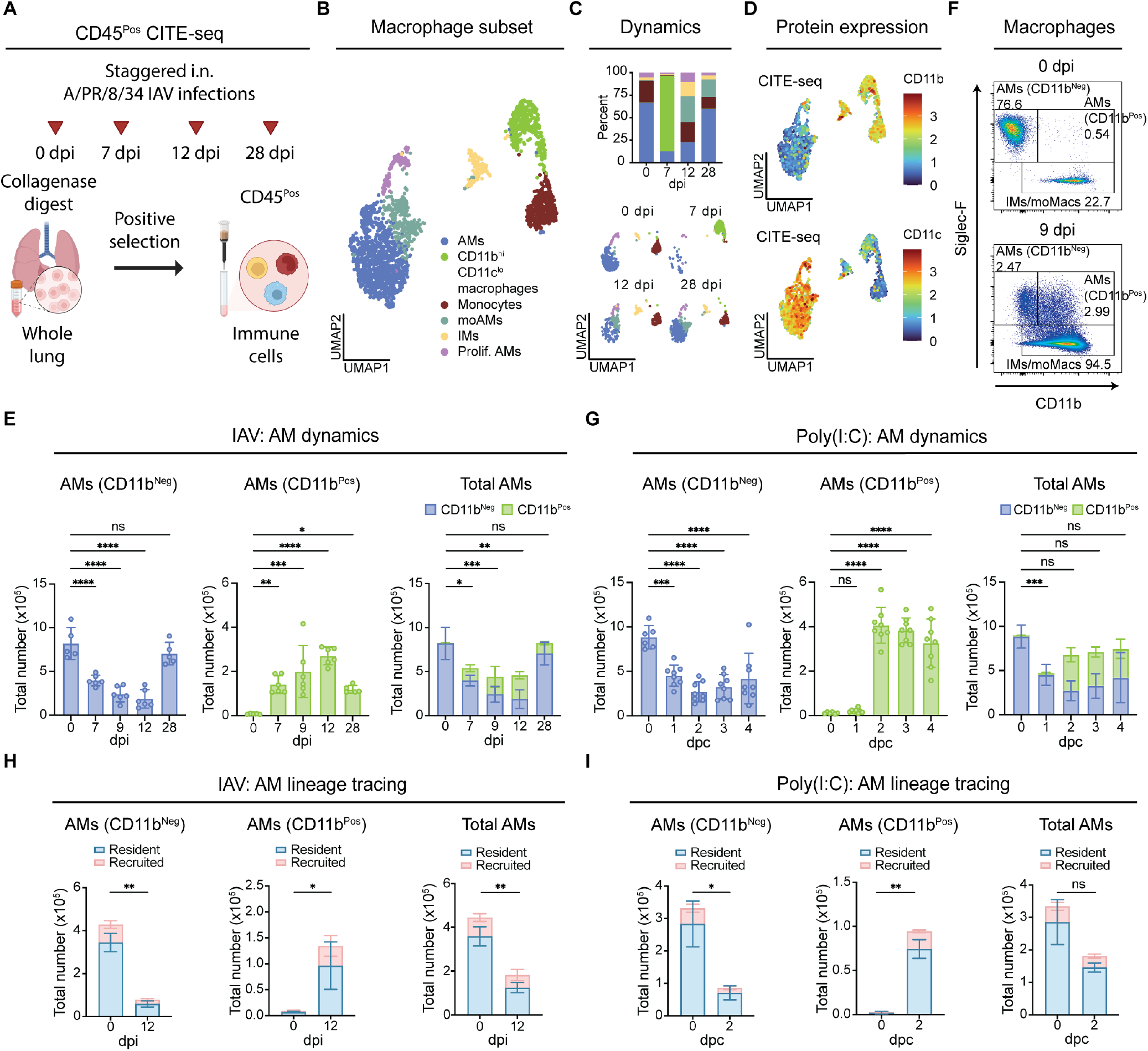
Exposure to viral stimuli induces a transient tissue resident-derived CD11b^Pos^ AM population. **(A–D)** CITE-seq characterization of myeloid cells during sublethal i.n. IAV infection (A/PR/8/34; 6.5 TCID_50_ for females and 32 TCID_50_ for males). Mice were infected and harvested at 0 (mock-infected), 7, 12, or 28 dpi (A; n = 1 male and 1 female mouse per group). UMAP of monocyte and macrophage subset (2,168 cells; B). Dynamics of macrophages over time (C). CD11b and CD11c protein expression plots (D). **(E–F)** Flow cytometry quantification of CD11b^Neg^, CD11b^Pos^, and total alveolar macrophages (AMs) after mock (0) or IAV infection (5 TCID_50_ ; 7, 9, 12, or 28 dpi; n = 5 - 6 mice per group) (E). Representative gating of CD11b^Neg^ and CD11b^Pos^ AMs, and IMs/monocyte-derived macrophages (moMacs) at 0 or 9 dpi (F). (**G**) Flow cytometry quantification of CD11b^Neg^, CD11b^Pos^, and total AMs during 0 (mock), 1, 2, 3, and 4 dpc with poly(I:C) (33.75 μg; n = 6 - 8 mice per group) pre-gated on live, CD45^Pos^, F4/80^Pos^, CD64^Pos^ singlets. **(H–I)** Flow cytometry quantification of *Ms4a3*^*Cre*^ LSL-tdTomato lineage tracing after viral challenge pregated on live, im. Resident tdTomato^Neg^ in blue and recruited (GMP-derived) tdTomato^Pos^ in red. Lineage tracing at 0 or 12 dpi (IAV; 5 TCID_50_ ; n = 3-4 mice per group) of CD11b^Neg^, CD11b^Pos^, and total AMs (H). Lineage tracing at 0 or 2 dpc (poly(I:C); 33.75 μg; n = 3 mice per group) of CD11b^Neg^, CD11b^Pos^, and total AMs (I). Data are representative of at least two independent experiments for (E, H, and I). Data from two independent experiments were pooled for (G). One-way ANOVA with Dunnet’s post-test for (E; Calculated on the sum of CD11b^Neg^ and CD11b^Pos^AMs for Total AMs), (G; Calculated on the sum of CD11b^Neg^ and CD11b^Pos^ AMs for Total AMs). T-tests for (H) and (I) calculated on the sum of tdTomato^Neg^ and tdTomato^Pos^ AMs. Error bars indicate standard deviation (SD). \**p* ≤ 0.05, * * *p* ≤ 0.005, * * \**p* ≤ 0.0005, * * * * *p* ≤ 0.0001.

To validate these scRNA-seq observations, we performed flow cytometry analysis following IAV infection (fig. S3D). Infection resulted in accumulation of neutrophils and monocytes, and an increase in IMs/moMacs (fig. S3E). We also observed the previously reported AM disappearance reaction (*15, 33*) at 7 dpi and throughout the course of infection (Fig. 2E). Concurrently, we observed the emergence of a CD11b^Pos^ CD11c^Pos^ Siglec-F^Pos^ macrophage population (Fig. 2E and 2F, and fig. S3F) likely representing the transient CD11b^Hi^ CD11c^Lo^ macrophages observed via CITE-seq (Fig. 2D). Based on surface marker expression (fig. S3D and S3F), we termed these cells CD11b^Pos^ AMs. When CD11b^Neg^ and CD11b^Pos^ AMs were quantified together, AM disappearance appeared to be more modest than previous reports (Fig. 2E).

As with the epithelial compartment, we hypothesized that the phenotypic changes in myeloid cells were mediated, at least in part, by IFN-I. To test this, we again employed our single-dose poly(I:C) challenge model. Poly(I:C) challenge resulted in no significant change in neutrophil numbers, an acute loss of monocytes, and a simultaneous increase in IM/moMacs (fig. S3G) suggesting IFN-I-driven differentiation of monocytes into macrophages. Poly(I:C) was also sufficient to recapitulate the loss of CD11b^Neg^ AMs and induction of CD11b^Pos^ AMs observed during IAV infection (Fig. 2G). Combining CD11b^Pos^ and CD11b^Neg^ AMs into a single population for quantification (Fig. 2G) again suggested reduced AM disappearance compared with previous reports after viral challenge.

Prior studies have demonstrated key roles for both monocyte-derived (*12*) and tissue-resident (*15*) AMs during IAV infection, prompting us to define the ontogeny of the transient CD11b^Pos^ AM population. To this end, we employed the *Ms4a3*^*Cre*^ tdTomato lineage tracing model (*34*), in which all granulocyte-monocyte progenitor (GMP)derived cells are labeled tdTomato^Pos^. Following IAV infection, monocytes, neutrophils, and IMs/moMacs were primarily recruited (fig. S4A), whereas CD11b^Neg^ AMs were predominantly resident-derived (Fig. 2H), as anticipated. Unexpectedly, CD11b^Pos^AMs also originated largely from resident cells rather than from recruited GMP-derived cells (Fig. 2H). A similar pattern was observed following poly(I:C) challenge, with monocytes, neutrophils, and IMs/moMacs recruited (fig. S4B) and both AM populations resident-derived (Fig. 2I). These results demonstrate that inflammatory cues within the alveolar niche after viral challenge alter canonical AM surface phenotypes.

### IFN-I signaling induces proliferative Sca-1^Pos^ ATIIs and promotes CD11b^Pos^ AM generation

In order to directly test the role of IFN-I in shaping ATII and AM phenotypes, we employed IFNAR1-blocking antibodies during poly(I:C) challenge to inhibit IFN-I signaling (Fig. 3A). Local i.t. IFNAR1 blockade was sufficient to abrogate induction of the Sca-1^Pos^ state and dampen overall ATII proliferation after challenge (Fig. 3B). Blockade of IFN-I signaling did not result in significant loss of ATIIs (fig. S5A). Furthermore, the Sca-1^Pos^ ATIIs persisting after blockade remained more proliferative relative to their Sca-1^Neg^ counterparts (fig. S5B).

**Figure 3.**
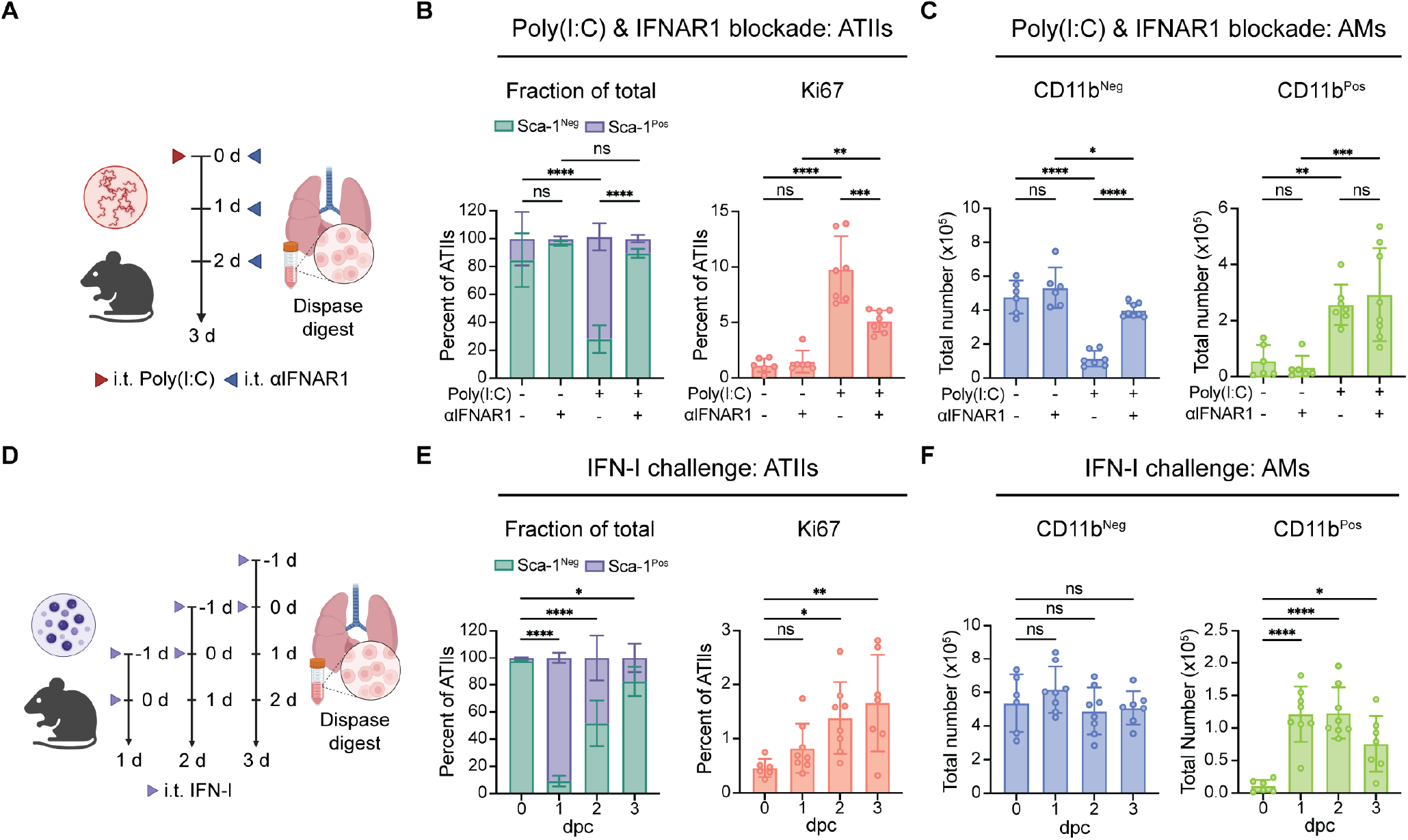
IFN-I signaling promotes induction of transient ISG^hi^ Sca-1^Pos^ ATIIs and CD11b^Pos^ AMs. (**A–C)** Flow cytometry quantification after IFNAR1 blockade (200 μg per treatment) and poly(I:C) challenge (33.75 μg) of epithelial and myeloid populations. Treatment and harvest scheme (A; n = 6 - 8 mice per group). Percent Sca-1^Neg/Pos^ATIIs of total or percent Ki67^Pos^ of ATIIs (B). CD11b^Neg^ and CD11b^Pos^ AM quantification (C). (**D–F**) Flow cytometry quantification after exogenous IFN-I challenge (i.t. 450 ng IFN-β per treatment). Treatment and harvest scheme (D; n = 6 - 8 mice per group). Percent Sca-1^Neg/Pos^ ATIIs of total or percent Ki67 ^Pos^ of ATIIs (E). CD11b^Neg^ and CD11b^Pos^ AM quantification (F). Data from two independent experiments were pooled for (B, C, E and F). One-way ANOVA with Dunnet’s post-test for (B), (C), (E), and (F). Calculated on Sca-1^Pos^ ATIIs for fraction of total plots. Error bars indicate standard deviation (SD). \**p* ≤ 0.05, * * *p* ≤ 0.005, * * \**p* ≤ 0.0005, * * * * *p* ≤ 0.0001.

Within the myeloid compartment, IFNAR1 blockade resulted in similar numbers of neutrophils (fig. S5C), monocytes (fig. S5D), and a significant decrease in IM/moMacs (fig. S5E). Blocking IFN-I signaling also resulted in a rescue of CD11b^Neg^ AM loss but did not affect induction of CD11b^Pos^ AMs following challenge (Fig. 3C and fig. S5F). These data suggest that IFN-I signaling via IFNAR1 contributes to AM loss but is not necessary for induction of the CD11b^Pos^state. This implies that alternative signals, such as IFN-II or IFN-III, may compensate to promote the CD11b^Pos^ AM state when IFN-I is blocked.

We next tested whether IFN-I was sufficient to induce our observed phenotypes within ATIIs and AMs (Fig. 3D). Local delivery of IFN-I was sufficient to induce both the Sca-1^Pos^ ATII state, as well as ATII proliferation, mirroring responses seen in IAV infection and poly(I:C) challenge (Fig. 3E, and fig. S5G, and S5H). Within the AM population, we did not observe a reduction in CD11b^Neg^ AMs but did detect an induction of CD11b^Pos^ AMs (Fig. 3F), without significant loss of total AMs (fig. S5I). These data suggest that IFN-I sensing is sufficient to induce CD11b expression in resident AMs. Exogenous IFN-I delivery reduced neutrophil and monocyte numbers (fig. S5J and S5K), while increasing IMs/moMac numbers (fig. S5L), further supporting a role for IFN-I in promoting monocyte to macrophage differentiation. Using the *Ms4a3*^Cre^ lineage tracer, we identified ontogeny consistent with that observed in IAV infection and poly(I:C) challenge for neutrophils, monocytes, IMs/moMacs, and AMs (fig. S5M-R). Taken together, these results suggest IFN-I signaling is necessary and sufficient to induce the Sca-1^Pos^ ATII state, and sufficient to promote the CD11b^Pos^ AM state.

### IFN-I priming mediates the ATII proliferative response via macrophage-derived OSM

A subset of tissue-resident AMs (trAMs) was retained and underwent phenotypic remodeling after viral challenge, raising the possibility that these cells are functionally important during inflammation. Specifically, CD11b^Pos^ AMs increased in number concurrently with Sca-1^Pos^ ATIIs (Fig. 1G, 1I, 2E, and 2G), suggesting a potential role for trAMs in driving ATII proliferation. To test this hypothesis, we developed a novel trAM depletion model. AMs rely on ATII-derived GM-CSF for survival (*35*). Accordingly, we delivered GM-CSF-blocking antibodies i.t. prior to challenge to deplete trAMs (fig. S6A) and confirmed depletion according to origin using *Ms4a3*^*Cre*^ lineage tracing mice. GM-CSF blockade led to a downward trend in neutrophil numbers after poly(I:C) treatment (fig. S6B) but had no effect on monocytes (fig. S6C) or IM/moMacs (fig. S6D). Importantly, GM-CSF blockade depleted both tissue-resident CD11b^Pos^ and CD11b^Neg^ AMs (Fig. 4A and fig. S6E). In the epithelial compartment, GM-CSF blockade did not significantly reduce total ATII numbers but did result in an accumulation of Sca-1^Pos^ ATIIs (Fig. 4B). Depletion of trAMs also attenuated ATII proliferation (Fig. 4B). These results implicate trAMs as a key source of proliferative signals and uncouple induction of the Sca-1^Pos^ ATII state from its proliferation.

**Figure 4.**
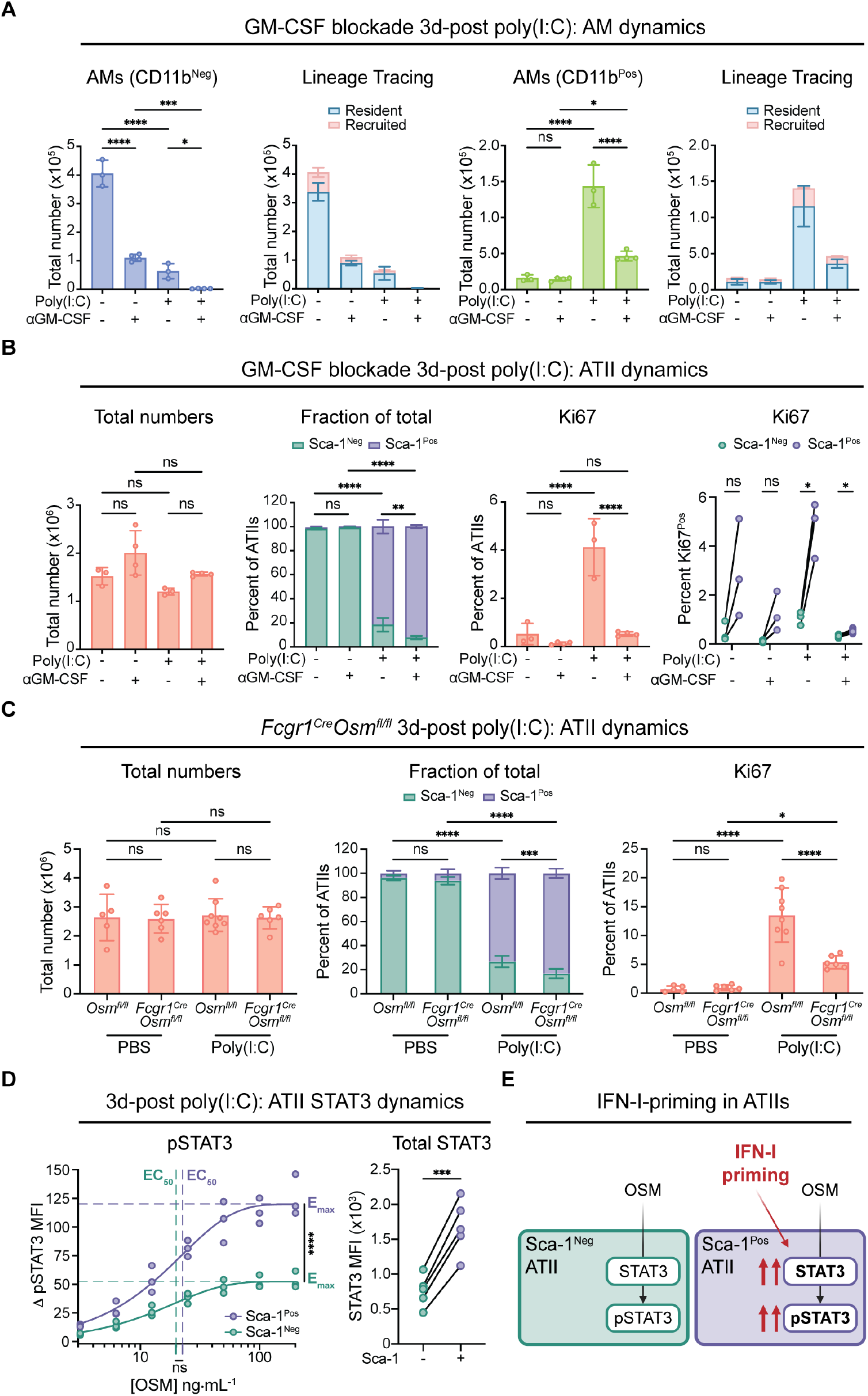
IFN-I priming enhances ATII responsiveness to macrophage-derived OSM. **(A–B)** Flow cytometry quantification of myeloid and epithelial populations during GM-CSF blockade (i.t. 1x daily for 3 days; days -3, -2, -1; 90 μg per treatment) and harvested at 3 d following i.t. poly(I:C) challenge (treated on day 0; i.t 33.75 μg) in *Ms4a3*^*Cre*^ tdTomato lineage tracing mice (n = 3 – 4 mice per group). Resident tdTomato^Neg^ in blue and recruited tdTomato^Pos^ in red. Quantification of CD11b^Neg^ and CD11b^Pos^ AM total numbers, and lineage deletion (A). Quantification of ATII total numbers, fraction of Sca-1^Neg/Pos^ ATIIs of total, percent Ki67^Pos^ of total ATIIs and percent Ki67^Pos^ of Sca-1^Neg/Pos^ ATIIs (B). **(C)** Flow cytometry quantification of ATII total numbers, fraction of Sca-1^Neg/Pos^ATIIs of total and percent Ki67^Pos^ of total ATIIs 3 d following i.t. poly(I:C) challenge (33.75 μg) in *Osm*^*fl/fl*^ or *Fcgr1*^*Cre*^*Osm*^*fl/fl*^ mice (n = 5 – 8 mice per group). **(D)** Flow cytometry quantification of Δ phospho-STAT3 mean fluorescence intensity (Δ pSTAT3 MFI) after poly(I:C) challenge (3 dpc; 33.75 μg) followed by *ex vivo* OSM stimulation of digested lung fractions across a range of doses (mock, 3.125, 6.25, 12.5, 25, 50, 100, and 200 ng*·*mL^-1^ with n = 3 biological replicates per OSM dose; calculated by subtracting out the average MFI of mock controls from the MFI of OSM stimulated samples) and total STAT3 MFI during poly(I:C) challenge (3 dpc; 33.75 μg; n = 5 mice per group; calculated by subtracting out fluorescence minus one controls from raw MFI). (**E**) Proposed model of IFN-I priming in ATIIs. Data are representative of at least two independent experiments for (A, B, and D). Data from two independent experiments were pooled for (C). One-way ANOVA with Dunnett’s post-test for (A), (B), and (C). Calculated on Sca-1^Pos^ATIIs for “Fraction of total” plots. Paired T-tests for (B; Sca-1^Neg^ vs. Sca-1^Pos^ percent Ki67^Pos^ plot) and (D) with Holm-Šídák correction for (B). A one-phase association model was used to draw lines of best-fit for the pSTAT3 dose response curves (D) and thereby also determined EC_50_ (tau) and E_max_ (plateau) values. To test if EC_50_ and E_max_ values were different, extra sum-of-squares F tests were performed on these values. Error bars indicate standard deviation (SD). \**p* ≤ 0.05, * * *p* ≤ 0.005, * * \**p* ≤ 0.0005, * * * * *p* ≤ 0.0001.

Previous work identified macrophage-derived oncostatin M (OSM) as a critical mediator of ATII proliferation, overcoming the anti-proliferative effects of IFN-I and restoring the epithelial barrier following viral-induced damage (*11*). Thus, we hypothesized that OSM may be the AM-derived signal mediating Sca-1^Pos^ ATII proliferation. To test this, we challenged *Fcgr1*^*Cre*^*Osm*^*fl/fl*^ mice, in which *Osm* is deleted specifically in macrophages (*11*), with poly(I:C). We found no significant differences in total ATII numbers but observed an accumulation Sca-1^Pos^ ATIIs following challenge (Fig. 4C). ATII proliferation was also reduced in mice lacking macrophage-derived OSM, indicating that OSM is a crucial mediator of IFN-primed Sca-1^Pos^ ATII proliferation (Fig. 4C).

We next sought to define the molecular mechanism underlying the enhanced proliferative response of Sca1^Pos^ ATIIs to OSM. In mice, OSM signals primarily through the heterodimeric receptor composed of gp130 and OSMR, activating downstream signaling pathways including STAT3 phosphorylation (pSTAT3) (*36, 37*). Given that loss of macrophage-derived OSM dampened Sca1^Pos^ ATII proliferation (Fig. 4C), we hypothesized that Sca-1^Pos^ ATIIs may have a greater potential to respond to OSM than their Sca-1^Neg^ counterparts. To interrogate this, we challenged mice with poly(I:C), stimulated lung suspensions *ex vivo* with increasing doses of OSM, and measured pSTAT3 (fig. S7A-C). Sca-1^Pos^ ATIIs exhibited a greater maximal response compared with Sca-1^Neg^ ATIIs (E_max_; Fig. 4D and fig. S7D), whereas the sensitivity to OSM remained unchanged between populations (EC_50_; Fig. 4D and fig. S7D). These results suggest that IFN-experienced ATIIs display enhanced signaling capacity without altered ligand sensitivity, consistent with stoichiometric scaling of the OSM response. Given this observation, and that *Stat3* is itself an ISG (*27*), we hypothesized that IFN-I signaling may increase STAT3 abundance in Sca-1^Pos^ ATIIs, thereby explaining their increased capacity to respond to OSM. Indeed, we found that Sca-1^Pos^ ATIIs expressed higher levels of total STAT3 compared with Sca-1^Neg^ATIIs (Fig. 4D, fig. S7D and S7E). Collectively, these observations suggest that IFN-I signaling functions beyond antiviral defense to prime alveolar epithelial cells for rapid renewal driven by tissue-resident macrophage-derived OSM (Fig. 4E and fig. S7F).

## Discussion

Our study supports a model in which IFN-I signaling prepares the lung for epithelial renewal following inflammatory challenge. We demonstrate that IFN-I contributes to the induction of an ISG^hi^ population of ATIIs marked by Sca-1 expression. Sca-1^Pos^ ATIIs exhibit enhanced proliferative capacity and organoid-forming efficiency relative to their Sca-1^Neg^ counterparts. Following viral challenge, proliferation of Sca-1^Pos^ ATIIs requires AM-derived OSM. Finally, we demonstrate that Sca-1^Pos^ ATIIs exhibit an enhanced responsiveness to OSM relative to Sca-1^Neg^ ATIIs due to heightened signaling capacity. Together, these findings reveal that viral stimuli induce coordinated IFN-I-dependent epithelial and macrophage states that prime the lung for renewal. In this context, IFN-I acts not only as a key antiviral defense mechanism but as a tuning signal that expands the capacity of the epithelium to integrate proliferative cues required for efficient repair.

Epithelial-intrinsic repair mechanisms in the lung have been well characterized, with one key route being ATII self-renewal and differentiation into ATIs following injury (*2*). Wnt signaling has been implicated in promoting ATII expansion (*38–41*), and several studies demonstrate that ATII-to-ATI differentiation occurs through transient intermediate states (*25, 26, 29*). Our data extend these models by positioning IFN-I priming as an upstream event that precedes previously described transitional states. We find that ISG^hi^ Sca-1^Pos^ ATIIs are transcriptionally similar to previously described activated/primed ATIIs and temporally precede their emergence. These observations suggest that IFN-I exposure may act as a calibration mechanism that enhances ATII capacity to engage canonical regenerative programs in response to subsequent signals. Consistent with this model, recent work has shown that permitting early inflammatory responses to proceed, followed by antiviral therapy and IFNAR blockade, reduces mortality following IAV infection (*42*).

Prior studies suggest that IFN-mediated priming may not be restricted to antiviral or IFN-I responses. IFN-I sensing by ATIIs has been shown to be required for preventing epithelial cell death and restoring barrier integrity following *Streptococcus pneumoniae* infection and HCl-induced injury (*43*). Similarly, a potentially analogous ATII population has been implicated in epithelial repair following *Pseudomonas aeruginosa* infection (*44, 45*). In addition, IFN-γ has been observed to promote proliferation in human lung organoids (*46*). These studies suggest that similar ATII states may be induced across both viral and bacterial contexts. Evidence from other tissues further supports a broader role for IFN in priming epithelial regeneration. In the gastrointestinal tract, mice deficient in both IFN-I and IFN-III signaling exhibit reduced proliferative responses within intestinal crypts following dextran sulfate sodium (DSS)-induced injury (*47*). Subsequent work demonstrated that blockade of IFN-III, but not IFNI alone, reduced proliferative responses after DSS injury (*48*), suggesting a possible combinatorial effect of IFN-I and IFN-III. Additionally, exogenous IFN-I has been shown to boost proliferation following radiation-induced damage (*49*), while endogenous IFN-γ promotes crypt proliferation, an effect dependent on intact IFN-γ signaling (*50*). IFNγ responses have also been linked to the emergence of a fetal-like Sca-1^Pos^ epithelial population (*51*), and similar IFN-associated epithelial states have been reported in other tissues such as the salivary gland (*52*) and prostate (*53*). Together, these studies point to a potentially conserved role for IFN-mediated epithelial priming across barrier tissues.

The presence of a minor Sca-1^Pos^ ATII population at steady state raises questions about a possible homeostatic role for IFN signaling in epithelial maintenance. Indeed, tonic IFN signaling has been previously reported in the homeostatic lung (*54*). Consistently, we found that Sca-1^Pos^ ATIIs present at baseline exhibit similar proliferative capacity and OSM sensitivity as their IAV or poly(I:C)induced counterparts. Supporting this idea, mice deficient in IFN signaling display reduced ATII cell turnover (*55*), and multiple tissue stem cell populations express ISGs (*56*). Interestingly, Sca-1-positivity and high STAT3 activity, both indicative of potential IFN-priming, have been associated with stemness and proliferation in models of IAV (*42*) and lung cancer (*57, 58*). Future studies will be required to define the source and functional role of tonic IFN in the maintenance of lung epithelial progenitor populations.

The role of macrophages in lung repair has recently come into sharper focus, yet the importance of ontogeny has not been fully defined. Several studies have implicated moAMs as mediators of repair following IAV infection (*12*), chemical damage (*29, 59*), and physical injury (*9*). Additional work has suggested that moAMs are drivers of pulmonary fibrosis (*60–63*). Macrophages of fetal origin have also been proposed to support tissue repair (*64*), and higher proportions of trAMs correlate with improved outcomes following IAV infection (*15*). Our GM-CSF blockade and *Ms4a3*^*Cre*^ tdTomato lineage-tracing data suggest a model in which trAMs undergo phenotypic remodeling and upregulate markers typically associated with monocyte-derived populations in response to inflammatory stimuli. These trAMs then promote ATII proliferation through OSM production. Together, our findings suggest that trAMs can adopt inflammation-associated states that actively drive epithelial regeneration.

The macrophage disappearance reaction is a phenomenon observed across a variety of tissues and inflammatory contexts, characterized by the apparent loss of macrophages during inflammation (*65*). Initially thought to reflect cell death or migration, subsequent work has revealed more complex processes. For instance, in the peritoneal cavity, macrophage disappearance was found to result from cellular adhesion and coagulation that leads to their absence from peritoneal fluid rather than cell loss (*66*). Similarly, a study in the lung demonstrated that trAMs remain physically present but are immobilized following IAV infection (*67*), suggesting that prior reports of AM loss may also reflect adhesion rather than depletion. Using comprehensive tissue digestion, a broad panel of surface markers, and GMP lineage tracing, we found that while some numerical loss occurs, a portion of the disappearance reflects phenotypic remodeling, including upregulation of the integrin CD11b. In IAV infection, poly(I:C) challenge, and IFN-I administration, we observe the emergence of a CD11b^Pos^ trAM cell state, revealing inflammation-driven macrophage plasticity.

Barrier tissues such as the lung are constantly exposed to potentially harmful stimuli, including viral infection. These encounters induce robust IFN-I responses that have previously been viewed as incompatible with tissue regeneration. Our findings challenge this paradigm by suggesting that IFN-I signaling promotes epithelial and macrophage states that actively facilitate repair. We propose that IFN-I may have evolved not only to control viral replication but also to initiate regenerative programs in parallel, enabling rapid restoration of tissue integrity after infection.

## Materials and Methods

### Mice

All animal experiments were performed in accordance with institutional regulations after protocol review and approval by the Institutional Animal Care and Use Committee (IACUC) at Harvard Medical School. Mice were bred and housed in a specific pathogen-free (SPF) facility with room temperature set to 71°F (+/-3°), humidity set to 50% (+/-15%), 12:12-hour light/dark cycle, and ad libitum access to food (LabDiet 5053) and water. C57BL/6J mice (RRID:IMSR_JAX:000664), *Ms4a3*^*Cre*^ (RRID:IMSR_JAX:036382), LSL-tdTomato (RRID:IMSR_JAX:007914), and β-act-EGFP (RRID:IMSR_JAX:006567) were obtained from Jackson Laboratories. *Fcgr1*^*Cre*^ mice were provided by Dr. Ming Li (MSKCC)^68^. *Osm*^*fl/fl*^ mice were bred to *Fcgr1*^*Cre*^ mice to achieve conditional deletion (*11*). Animals were euthanized by administration of a lethal dose of ketamine/xylazine mixture. Female mice aged 8 to 12 weeks were used for all experiments, unless otherwise specified.

### Infection and *in vivo* treatments

Mice were acclimated in our mouse facility for at least one week prior to experimentation. All IAV infections were performed using the A/Puerto Rico/8/1934 (H1N1) strain (Charles River Laboratories). Infectious dose was determined via 50% tissue culture infectious dose (TCID_50_) as previously described (*69*) In short, MDCKs were infected with serial dilutions of IAV for 48hrs at 35 °C. Supernatant containing virus was mixed with chicken red blood cells (Rockland) to perform a hemagglutination assay and the TCID_50_ was then calculated using the Reed-Muench method. For IAV or mock infection, mice were anesthetized with ketamine/xylazine and inoculated intranasally with 5-32 TCID_50_ IAV or PBS in a 30 μL volume. For i.t. treatments, mice were anesthetized with isoflurane/propanediol. Mice received i.t. treatments of high–molecular weight poly(I:C) (33.75 μg; InvivoGen), exogenous IFN-I (450-700 ng IFN-β; BioLegend), or blocking antibodies against IFNAR1 (200 μg per treatment; InVivoPlus anti-mouse IFNAR1, Bio X Cell) or GM-CSF (50 μg per treatment; InVivoPlus anti-mouse GM-CSF, Bio X Cell), with corresponding rat IgG1a (αIFNAR1) or IgG2a (GM-CSF) isotype controls, each diluted in PBS or pH-matched buffer to a final volume of 50–54 μLs.

### Lung isolation for flow cytometry, single cell RNA-seq, and cell sorting

Lungs were harvested and processed as previously described^70^. Briefly, mice were euthanized and perfused with PBS containing 2 mM EDTA. For all analyses, the trachea was exposed and cannulated with a 22-gauge catheter to lavage the lungs with PBS for bronchoalveolar lavage fluid (BALF) collection, followed by inflation with dispase (Corning; 50 caseinolytic units/mL). Lungs were removed, lobes were finely minced, and tissue was digested in RPMI containing DNase (Sigma;100 μg/mL) and Liberase (Sigma; 83 μg/mL) at 37 °C for 40 min with agitation. BALF and lung digests were filtered through 70 μm strainers to generate single-cell suspensions and treated with ACK lysis buffer (Thermo Fisher Scientific) to remove red blood cells. Cell viability was assessed using Zombie Aqua (1:150; BioLegend) for flow cytometry and DAPI (10.9 μM; BioLegend) for cell sorting. Cells were Fc-blocked with anti-mouse CD16/32 (1:250) prior to staining. For surface and intracellular staining, cells were fixed and permeabilized using Foxp3/Transcription factor Fix/Perm (eBioscience) and stained for 1 h at room temperature. For EdU analyses, cells were surface stained, fixed in 4% PFA for 15 min, and EdU incorporation was detected using the Click-iT Plus EdU assay (Thermo Fisher) according to the manufacturer’s instructions. For STAT3 and pSTAT3 staining, cells were permeabilized overnight in 90% methanol at 20 °C prior to staining. Data were acquired on a BD Symphony A5 or A1 and analyzed using FlowJo (10.10.1; Waters Bioscience); cell sorting was performed on a BD Aria 561. Absolute cell numbers were calculated using 123count eBeads (Thermo Fisher).

### scRNA-seq and CITE-seq sample collection

8-week-old male and female C57BL/6J mice were infected with 6.5 TCID_50_ for females and 32 TCID_50_ for males at staggered time points to synchronize tissue harvests across two consecutive days, with one 10x Genomics encapsulation performed per day. On one day, a collagenase-based digest was used to preserve immune cell surface epitopes, followed by CD45 magnetic bead enrichment for CITE-seq. On the other day, a dispase-based digest was used to isolate epithelial and stromal cells, followed by CD45 and CD31 magnetic bead depletion for scRNA-seq. For each encapsulation, lungs from male and female mice at each time point were processed in parallel. Following enzymatic digestion and filtration (as above), cells were labeled with sample-specific TotalSeq™-B0301B0309 anti-mouse Hashtag antibodies (BioLegend) and incubated with anti-CD45 microbeads (day 1; Miltenyi) or biotinylated anti-CD45 (1:200; BioLegend) and anti-CD31 (1:200; BioLegend) antibodies followed by streptavidin-conjugated magnetic beads (day 2; Miltenyi). Magnetic separations were performed using MS columns (Miltenyi) on a MACS separator (Miltenyi), collecting either the positively selected (immune cell) or flow-through (epithelial and stromal cell) fractions. Cells were counted, assessed for viability, pooled to equal representation across samples, and adjusted to a final input of 50,000 cells per encapsulation prior to 10x Genomics library preparation.

### Upstream processing of 10x single-cell and CITE-seq data

For single-cell RNAseq, raw 10X BCL files are demultiplexed into FASTQ files using the Cell Ranger function cellranger mkfastq. The CD45 enriched library had GEX index SI-TT-A1 and FBC index SI-NT-A2. The Epithelial and stromal library had GEX index SI-TT-G12 and FBC index SI-NT-D1. Each library was split across two lanes of 10x in sequencing. Count matrices for GEX and FBC data for both libraries are generated using the function cellranger count. GRCm38 release M25 appended with the A/Puerto Rico/8/1934 genome (GCF_000865725.1) was used in addition the following versions of software: Cell Ranger 8.0.1 & R 4.1.2.

### Downstream analysis of scRNA-seq and CITE-seq data

Epithelial and stromal cell single-cell RNA-seq and immune cell CITE-seq datasets were analyzed in R (4.4.1) using Seurat (5.3.1). Cell Ranger outputs were loaded using Seurat and hashtag oligonucleotide (HTO) classifications were used to demultiplex samples using HTOde-mux() function from Seurat. HTO identities were used to annotate sex and IAV infection timepoint, and these annotations were added as metadata fields. Time points were ordered explicitly for downstream analyses. Doublets and contaminating immune or stromal populations (respectively) were excluded. For the immune population, the following QC cutoffs were used; RNA counts per cell 500, ADT counts per cell > 147, Genes per cell > 300, percent mitochondrial reads < 5%. For the stromal population, the following QC cutoffs were used; nCounts_RNA > 550, nCounts_ADT > 0 (no protein data), nFeatures_RNA 500, percent_mito < 5%. Data underwent conventional normalization (NormalizeData(“LogNormalize”)), variable feature selection (FindVariableFeatures(2000)), scaling (ScaleData()), PCA using (runPCA()), graph-based clustering (findNeighbors(), findClusters(), runUMAP()) using Seurat’s pipeline. Major clusters were annotated based on canonical marker expression and differential gene analysis, and annotations were stored as metadata for downstream analyses. Subsets of interest (ATII and monocyte/macrophage populations) were reprocessed independently as above with higher-resolution clustering, and bespoke gene set enrichment analysis. Gene set enrichment (ISG, ADI, and activated ATIIs), was quantified using module scoring and visualized using Nebulosa (1.14.0) density plots and were also used for subcluster annotation.

### Organoid culture

β-act-EGFP mice were processed as above to obtain a single-cell suspension. FACS-isolated mouse lung eGFP^Pos^, viable, Sca-1^Neg^ or Sca-1^Pos^ ATIIs were resuspended in 3D media (DMEM/F12 supplemented with 10% FBS, penicillin/streptomycin, 1 mM HEPES, and insulin/transferrin/selenium (Corning)) at a concentration of 5,000 live cells (trypan blue negative) per 50 μL. As supporting cells, a mix of neonatal stromal cells were isolated as described (*32*). The stromal cells were pelleted and resuspended in growth factor reduced (GFR) Matrigel (Corning) at a concentration of 50,000 cells per 50 μL. Equal volumes of cells in 3D media and supporting cells in GFR Matrigel were mixed and 100 μL were pipetted into a Transwell (Corning). Plates were incubated for 20 minutes at 37 °C, 5% CO2 until Matrigel solidified. Finally, 500 μL of 3D media was added to the bottom of the well. 3D media was changed every other day.

### Organoid imaging and analysis

Live-cell imaging was performed using an In Cell Analyzer 6000 equipped with a sCMOS camera. Images were acquired using a Plan Apo 4X objective (NA 0.20). To visualize eGFP organoids, a 488 nm laser was utilized with a 525/20 nm emission filter. Images were generated as maximum intensity projection of 51 Z-slices (step size of 50 μm). Raw images were stitched using a cross-correlation scheme and enhanced using non-local mean filtering. Automated organoid detection and segmentation were performed using a custom template-matching and morphological analysis pipeline in MATLAB. To identify putative centers, Gaussian-blurred stitched grayscale images were analyzed via 2D normalized cross-correlation against a filter bank of solid and edge-emphasized disk templates of varying radii. The resulting maximum correlation maps were adaptively thresholded to generate initial binary masks. Overlapping structures were computationally separated by analyzing object eccentricity and bisecting boundary “bottlenecks.” Finally, organoids falling below a 5000 μm^2^ area threshold (80 μm diameter) were discarded. The custom scripts developed for this analysis are available at https://doi.org/10.5281/zenodo.18750772. Organoid forming efficiency (OFE) was calculated by dividing the total number of organoids from the automated counts from each well and dividing by the total number of live cells seeded within each well (5,000 seeded cells).

### Data analysis and statistics

Statistical tests were performed using GraphPad Prism (10.6.1), with significance defined as *p* ≤ 0.05. One-way ANOVA was applied to compare the means across more than two independent groups. For post-hoc comparison of selected groups means we performed Dunnett’s test. For two group comparisons within the same mouse a paired *t* -test was used. For two-group comparisons across different mice a student *t* -test was used. For dose-response curves a one-phase association model was performed to draw lines of best-fit and determine EC_50_ and E_max_. To test if EC_50_ and E_max_ values were different we performed extra sum-of-squares F tests on these values.

## Acknowledgments

We thank all current and past members of the Franklin laboratory for their support and helpful discussions. We thank M. Henderson, O. Casey, and I. McGill for their assistance with scRNA-seq and CITE-seq. We also thank the HMS Immunology Flow Cytometry Core, Microscopy Resources on the North Quad (MicRoN) Core, and the Harvard Center for Comparative Medicine, especially C. Araneo, J. Giaquinto, G. Haskett, and P.V. Anekal. We thank A. Wang and R. Medzhitov for providing feedback on the manuscript and R. Jackson for experimental discussions. Figure schematics were created using BioRender.

## Funding

National Institutes of Health grant R35GM150816 (R.A.F.)

National Institutes of Health grant F31HL172650 (A.Y.B.V)

National Institutes of Health grant TL1TR002543 (Y.L., M.A.C., P.R-M.)

Harvard Stem Cell Institute (R.A.F.)

Harvard FAS Dean’s Competitive Fund for Promising Scholarship (R.A.F.)

Charles H. Hood Foundation (R.A.F.)

National Science Foundation DGE1122492 (P.R-M.)

Gene Lay Institute of Immunology and Inflammation Postdoctoral Fellowship (A.O.M.)

## Author contributions

Conceptualization: A.Y.B.V., D.A.H., A.O.M., R.A.F.

Formal analysis: A.Y.B.V., L.T.

Funding acquisition: R.A.F.

Investigation: A.Y.B.V., D.A.H., A.O.M., Y.L., S.M.D., M.H., L.T., S.B., M.A.C., P.R-M., A.L., I.B., D.S., A.S.

Methodology: D.A.H., A.O.M, F.C., C.B., C.F.K., R.A.F.

Supervision: F.C., C.F.K., R.A.F.

Visualization: A.Y.B.V., R.A.F.

Writing—Original Draft: A.Y.B.V., R.A.F.

Writing—Review and Editing: A.Y.B.V., D.A.H., A.O.M., Y.L., M.H., M.A.C., C.F.K., R.A.F.

## Data and materials availability

Requests for materials should be directed to the corresponding author. Sequencing data are available upon request, and will be released publicly upon publication.

## Bibliography

1. X. Wei, H. Narasimhan, B. Zhu, J. Sun, Host Recovery from Respiratory Viral Infection. Annu Rev Immunol 41, 277–300 (2023).

2. M. C. Basil, J. Katzen, A. E. Engler, M. Guo, M. J. Herriges, J. J. Kathiriya, R. Windmueller, A. B. Ysasi, W. J. Zacharias, H. A. Chapman, D. N. Kotton, J. R. Rock, H. W. Snoeck, G. Vunjak-Novakovic, J. A. Whitsett, E. E. Morrisey, The Cellular and Physiological Basis for Lung Repair and Regeneration: Past, Present, and Future. Cell Stem Cell 26, 482–502 (2020).

3. Z. Lv, Z. Liu, K. Liu, X. Lin, W. Pu, Y. Li, H. Zhao, Y. Xi, P. Sui, A. E. Vaughan, A. Gillich, B. Zhou, Alveolar regeneration by airway secretory-cell-derived p63+ progenitors. Cell Stem Cell 31, 1685-1700.e6 (2024).

4. D. L. Jones, M. P. Morley, X. Li, Y. Ying, G. Zhao, S. E. Schaefer, L. R. Rodriguez, F. L. Cardenas-Diaz, S. Li, S. Zhou, U. V. Chembazhi, M. Kim, C. Shen, A. Nottingham, S. M. Lin, E. Cantu, J. M. Diamond, M. C. Basil, A. E. Vaughan, E. E. Morrisey, An injury-induced mesenchymal-epithelial cell niche coordinates regenerative responses in the lung. Science 386, eado5561 (2024).

5. K. Liu, X. Meng, Z. Liu, M. Tang, Z. Lv, X. Huang, H. Jin, X. Han, X. Liu, W. Pu, H. Zhu, B. Zhou, Tracing the origin of alveolar stem cells in lung repair and regeneration. Cell 187, 2428-2445.e20 (2024).

6. C. F. B. Kim, E. L. Jackson, A. E. Woolfenden, S. Lawrence, I. Babar, S. Vogel, D. Crowley, R. T. Bronson, T. Jacks, Identification of Bronchioalveolar Stem Cells in Normal Lung and Lung Cancer. Cell 121, 823–835 (2005).

7. Q. Liu, K. Liu, G. Cui, X. Huang, S. Yao, W. Guo, Z. Qin, Y. Li, R. Yang, W. Pu, L. Zhang, L. He, H. Zhao, W. Yu, M. Tang, X. Tian, D. Cai, Y. Nie, S. Hu, T. Ren, Z. Qiao, H. Huang, Y. A. Zeng, N. Jing, G. Peng, H. Ji, B. Zhou, Lung regeneration by multipotent stem cells residing at the bronchioalveolar-duct junction. Nat. Genet. 51, 728–738 (2019).

8. H. Aegerter, B. N. Lambrecht, C. V. Jakubzick, Biology of lung macrophages in health and disease. Immunity 55, 1564–1580 (2022).

9. A. J. Lechner, I. H. Driver, J. Lee, C. M. Conroy, A. Nagle, R. M. Locksley, J. R. Rock, Recruited Monocytes and Type 2 Immunity Promote Lung Regeneration following Pneumonectomy. Cell Stem Cell 21, 120-134.e7 (2017).

10. I. G. Wong, M. Paschini, J. Stark, A. B. Vazquez, V. Ya, A. L. Moye, S. M. Dang, M. F. Trovero, E. L. Thompson, S. Ahmed, F. N. Chaudhry, A. Shehaj, M. J. Rouhani, R. Bronson, S. M. Janes, S. P. Rowbotham, J. A. Zepp, R. A. Franklin, C. F. Kim, Airway injury induces alveolar epithelial responses mediated by macrophages. Cell Rep. 45, 116881 (2026).

11. D. A. Hoagland, P. Rodríguez-Morales, A. O. Mann, Y. B. Vazquez, S. Yu, A. Lai, H. Kane, S. M. Dang, Y. Lin, L. Thorens, S. Begum, M. A. Castro, S. D. Pope, J. Lim, S. Li, X. Zhang, M. O. Li, C. F. Kim, R. Jackson, R. Medzhitov, R. A. Franklin, Macrophage-derived oncostatin M repairs the lung epithelial barrier during inflammatory damage. Science 389, 169–175 (2025).

12. C. Ruscitti, J. Abinet, P. Maréchal, M. Meunier, C. de Meeûs, D. Vanneste, P. Janssen, M. Dourcy, M. Thiry, F. Bureau, C. Schneider, B. Machiels, A. Hidalgo, F. Ginhoux, G. Dewals, J. Guiot, F. Schleich, M.-M. Garigliany, A. Bellahcène, C. Radermecker, T. Marichal, Recruited atypical Ly6G+ macrophages license alveolar regeneration after lung injury. Sci. Immunol. 9, eado1227–eado1227 (2024).

13. B. B. Ural, S. T. Yeung, P. Damani-Yokota, J. C. Devlin, M. de Vries, P. Vera-Licona, T. Samji, C. M. Sawai, G. Jang, O. A. Perez, Q. Pham, L. Maher, P. Loke, M. Dittmann, B. Reizis, K. M. Khanna, Identification of a nerve-associated, lung-resident interstitial macrophage subset with distinct localization and immunoregulatory properties. Science Immunology 5 (2020).

14. L. Y. Hung, D. Sen, T. K. Oniskey, J. Katzen, N. A. Cohen, A. E. Vaughan, W. Nieves, A. Urisman, M. F. Beers, M. F. Krummel, D. B. R. Herbert, Macrophages promote epithelial proliferation following infectious and noninfectious lung injury through a Trefoil factor 2-dependent mechanism. Mucosal Immunology 12, 64–76 (2019).

15. F. Li, F. Piattini, L. Pohlmeier, Q. Feng, H. Rehrauer, M. Kopf, Monocyte-derived alveolar macrophages autonomously determine severe outcome of respiratory viral infection. Sci Immunol 7, eabj5761 (2022).

16. Y. Kumagai, O. Takeuchi, H. Kato, H. Kumar, K. Matsui, E. Morii, K. Aozasa, T. Kawai, S. Akira, Alveolar Macrophages Are the Primary Interferon-α Producer in Pulmonary Infection with RNA Viruses. Immunity 27, 240–252 (2007).

17. H. M. Lazear, J. W. Schoggins, M. S. Diamond, Shared and Distinct Functions of Type I and Type III Interferons. Immunity 50, 907–923 (2019).

18. D. Boehmer, I. Zanoni, Interferons in health and disease. Cell 188, 4480–4504 (2025).

19. F. McNab, K. Mayer-Barber, A. Sher, A. Wack, A. O’Garra, Type I interferons in infectious disease. Nat Rev Immunol 15, 87–103 (2015).

20. B. Mishra, L. B. Ivashkiv, Interferons and epigenetic mechanisms in training, priming and tolerance of monocytes and hematopoietic progenitors. Immunol. Rev. 323, 257–275 (2024).

21. K. Högner, T. Wolff, S. Pleschka, S. Plog, A. D. Gruber, U. Kalinke, H.-D. Walmrath, J. Bodner, S. Gattenlöhner, P. Lewe-Schlosser, M. Matrosovich, W. Seeger, J. Lohmeyer, S. Herold, Macrophage-expressed IFN-β Contributes to Apoptotic Alveolar Epithelial Cell Injury in Severe Influenza Virus Pneumonia. Plos Pathog 9, e1003188 (2013).

22. H. Katsura, Y. Kobayashi, P. R. Tata, B. L. M. Hogan, IL-1 and TNFα Contribute to the Inflammatory Niche to Enhance Alveolar Regeneration. Stem Cell Rep 12, 657–666 (2019).

23. J. Major, S. Crotta, M. Llorian, T. M. McCabe, H. H. Gad, S. L. Priestnall, R. Hartmann, A. Wack, Type I and III interferons disrupt lung epithelial repair during recovery from viral infection. Sci New York N Y 369, 712–717 (2020).

24. A. Broggi, S. Ghosh, B. Sposito, R. Spreafico, F. Balzarini, A. L. Cascio, N. Clementi, M. D. Santis, N. Mancini, F. Granucci, I. Zanoni, Type III interferons disrupt the lung epithelial barrier upon viral recognition. Science 369, 706–712 (2020).

25. M. Strunz, L. M. Simon, M. Ansari, J. J. Kathiriya, I. Angelidis, C. H. Mayr, G. Tsidiridis, M. Lange, L. F. Mattner, M. Yee, P. Ogar, A. Sengupta, I. Kukhtevich, R. Schneider, Z. Zhao, C. Voss, T. Stoeger, J. H. L. Neumann, A. Hilgendorff, J. Behr, M. O’Reilly, M. Lehmann, G. Burgstaller, M. Königshoff, H. A. Chapman, F. J. Theis, H. B. Schiller, Alveolar regeneration through a Krt8+ transitional stem cell state that persists in human lung fibrosis. Nature Communications 11 (2020).

26. Y. Kobayashi, A. Tata, A. Konkimalla, H. Katsura, R. F. Lee, J. Ou, N. E. Banovich, J. A. Kropski, P. R. Tata, Persistence of a regeneration-associated, transitional alveolar epithelial cell state in pulmonary fibrosis. Nat. Cell Biol. 22, 934–946 (2020).

27. J. W. Schoggins, S. J. Wilson, M. Panis, M. Y. Murphy, C. T. Jones, P. Bieniasz, C. M. Rice, A diverse range of gene products are effectors of the type I interferon antiviral response. Nature 472, 481–485 (2011).

28. M. A. G. Essers, S. Offner, W. E. Blanco-Bose, Z. Waibler, U. Kalinke, M. A. Duchosal, A. Trumpp, IFNα activates dormant haematopoietic stem cells in vivo. Nature 458, 904–908 (2009).

29. J. Choi, J.-E. Park, G. Tsagkogeorga, M. Yanagita, B.-K. Koo, N. Han, J.-H. Lee, Inflammatory Signals Induce AT2 Cell-Derived Damage-Associated Transient Progenitors that Mediate Alveolar Regeneration. Cell Stem Cell 27, 366-382.e7 (2020).

30. H. Narasimhan, I. S. Cheon, W. Qian, S. S. Hu, T. Parimon, C. Li, N. Goplen, Y. Wu, X. Wei, Y. M. Son, E. Fink, G. de A. Santos, J. Tang, C. Yao, L. Muehling, G. Canderan, A. Kadl, A. Cannon, S. Young, R. Hannan, G. Bingham, M. Arish, A. S. Chaudhari, J. sub Im, C. L. R. Mattingly, P. Pramoonjago, A. Marchesvsky, J. Sturek, J. E. Kohlmeier, Y. M. Shim, J. Woodfolk, C. Zang, P. Chen, J. Sun, An aberrant immune–epithelial progenitor niche drives viral lung sequelae. Nature, 1–9 (2024).

31. H. Katsura, V. Sontake, A. Tata, Y. Kobayashi, C. E. Edwards, B. E. Heaton, A. Konkimalla, T. Asakura, Y. Mikami, E. J. Fritch, P. J. Lee, N. S. Heaton, R. C. Boucher, S. H. Randell, R. S. Baric, P. R. Tata, Human Lung Stem Cell-Based Alveolospheres Provide Insights into SARS-CoV-2-Mediated Interferon Responses and Pneumocyte Dysfunction. Cell Stem Cell 27, 890-904.e8 (2020).

32. J.-H. Lee, T. Tammela, M. Hofree, J. Choi, N. D. Marjanovic, S. Han, D. Canner, K. Wu, M. Paschini, D. H. Bhang, T. Jacks, A. Regev, C. F. Kim, Anatomically and Functionally Distinct Lung Mesenchymal Populations Marked by Lgr5 and Lgr6. Cell 170, 1149-1163.e12 (2017).

33. H. Aegerter, J. Kulikauskaite, S. Crotta, H. Patel, G. Kelly, E. M. Hessel, M. Mack, S. Beinke, A. Wack, Influenza-induced monocyte-derived alveolar macrophages confer prolonged antibacterial protection. Nature Immunology 21, 145–157 (2020).

34. Z. Liu, Y. Gu, S. Chakarov, C. Bleriot, I. Kwok, X. Chen, A. Shin, W. Huang, R. J. Dress, C. A. Dutertre, A. Schlitzer, J. Chen, L. G. Ng, H. Wang, Z. Liu, B. Su, F. Ginhoux, Fate Mapping via Ms4a3-Expression History Traces Monocyte-Derived Cells. Cell 178, 1509-1525.e19 (2019).

35. J. Gschwend, S. P. M. Sherman, F. Ridder, X. Feng, H.-E. Liang, R. M. Locksley, B. Becher, C. Schneider, Alveolar macrophages rely on GM-CSF from alveolar epithelial type 2 cells before and after birth. The Journal of experimental medicine 218, e20210745 (2021).

36. C. D. Richards, The enigmatic cytokine oncostatin m and roles in disease. ISRN Inflamm 2013, 512103 (2013).

37. H. M. Hermanns, Oncostatin M and interleukin-31: Cytokines, receptors, signal transduction and physiology. Cytokine Growth Factor Rev 26, 545–558 (2015).

38. W. J. Zacharias, D. B. Frank, J. A. Zepp, M. P. Morley, F. A. Alkhaleel, J. Kong, S. Zhou, E. Cantu, E. E. Morrisey, Regeneration of the lung alveolus by an evolutionarily conserved epithelial progenitor. Nature 555, 251–255 (2018).

39. D. B. Frank, T. Peng, J. A. Zepp, M. Snitow, T. L. Vincent, I. J. Penkala, Z. Cui, M. J. Herriges, M. P. Morley, S. Zhou, M. M. Lu, E. E. Morrisey, Emergence of a Wave of Wnt Signaling that Regulates Lung Alveologenesis by Controlling Epithelial Self-Renewal and Differentiation. Cell Rep. 17, 2312–2325 (2016).

40. A. Toth, P. Kannan, J. Snowball, M. Kofron, J. A. Wayman, J. P. Bridges, E. R. Miraldi, D. Swarr, W. J. Zacharias, Alveolar epithelial progenitor cells require Nkx2-1 to maintain progenitor-specific epigenomic state during lung homeostasis and regeneration. Nat. Commun. 14, 8452 (2023).

41. A. N. Nabhan, D. G. Brownfield, P. B. Harbury, M. A. Krasnow, T. J. Desai, Single-cell Wnt signaling niches maintain stemness of alveolar type 2 cells. Science 359, 1118–1123 (2018).

42. H. Ichise, E. Speranza, F. L. Russa, T. Z. Veres, C. J. Chu, A. Gola, B. H. Clark, R. N. Germain, Rebalancing viral and immune damage versus repair prevents death from lethal influenza infection. Science 390, eadr4635 (2025).

43. B. B. Maier, A. Hladik, K. Lakovits, A. Korosec, R. Martins, J. B. Kral, I. Mesteri, B. Strobl, M. Müller, U. Kalinke, M. Merad, S. Knapp, Type I interferon promotes alveolar epithelial type II cell survival during pulmonary Streptococcus pneumoniae infection and sterile lung injury in mice. Eur J Immunol 46, 2175–2186 (2016).

44. Y. Liu, V. S. Kumar, W. Zhang, J. Rehman, A. B. Malik, Activation of Type II Cells into Regenerative Stem Cell Antigen-1+ Cells during Alveolar Repair. Am. J. Respir. Cell Mol. Biol. 53, 113–124 (2015).

45. Y. Liu, R. T. Sadikot, G. R. Adami, V. V. Kalinichenko, S. Pendyala, V. Natarajan, Y. Zhao, A. B. Malik, FoxM1 mediates the progenitor function of type II epithelial cells in repairing alveolar injury induced by Pseudomonas aeruginosa. J. Exp. Med. 208, 1473–1484 (2011).

46. A. F. M. Dost, K. Balážová, C. P. Casellas, L. M. van Rooijen, W. Epskamp, G. J. F. van Son, W. J. van de Wetering, C. Lopez-Iglesias, H. Begthel, P. J. Peters, N. Smakman, J. H. van Es, H. Clevers, Interferon-γ selectively promotes survival of alveolar progenitor cells in a human lung organoid model. EMBO J., 1–32 (2026).

47. C. McElrath, V. Espinosa, J.-D. Lin, J. Peng, R. Sridhar, O. Dutta, H.-C. Tseng, S. V. Smirnov, H. Risman, M. J. Sandoval, V. Davra, Y.-J. Chang, B. P. Pollack, R. B. Birge, M. Galan, A. Rivera, J. E. Durbin, S. V. Kotenko, Critical role of interferons in gastrointestinal injury repair. Nat. Commun. 12, 2624 (2021).

48. K. K. Jena, J. Mambu, D. Boehmer, B. Sposito, V. Millet, J. de S. Casal, H. I. Muendlein, R. Spreafico, R. Fenouil, L. Spinelli, S. Wurbel, C. Riquier, F. Galland, P. Naquet, L. Chasson, M. Elkins, V. Mitsialis, N. Ketelut-Carneiro, K. B. Gwilt, J. R. Thiagarajah, H.-B. Ruan, Z. Lin, E. Lien, F. Shao, J. Chou, A. Poltorak, J. Ordovas-Montanes, K. A. Fitzgerald, S. B. Snapper, A. Broggi, I. Zanoni, Type III interferons induce pyroptosis in gut epithelial cells and impair mucosal repair. Cell 187, 7533-7550.e23 (2024).

49. B. J. Leibowitz, G. Zhao, L. Wei, H. Ruan, M. Epperly, L. Chen, X. Lu, J. S. Greenberger, L. Zhang, J. Yu, Interferon b drives intestinal regeneration after radiation. Sci. Adv. 7, eabi5253 (2021).

50. S. Takashima, R. Sharma, W. Chang, M. Calafiore, Y.-Y. Fu, S. A. Jansen, T. Ito, A. Egorova, J. Kuttiyara, V. Arnhold, J. Sharrock, E. Santosa, O. Chaudhary, H. Geiger, H. Iwasaki, C. Liu, J. Sun, N. Robine, L. Mazutis, C. A. Lindemans, A. M. Hanash, STAT1 regulates immune-mediated intestinal stem cell proliferation and epithelial regeneration. Nat. Commun. 16, 138 (2025).

51. Y. M. Nusse, A. K. Savage, P. Marangoni, A. K. M. Rosendahl-Huber, T. A. Landman, F. J. de Sauvage, R. M. Locksley, O. D. Klein, Parasitic helminths induce fetal-like reversion in the intestinal stem cell niche. Nature 559, 109–113 (2018).

52. D. Cinat, R. van der Wal, M. Baanstra, A. Soto-Gamez, R. Maturi, A. L. J. Bruin, U. Brouwer, M.-J. van Goethem, M. A. T. M. van Vugt, L. Barazzuol, R. P. Coppes, IFN-I signaling enhances salivary gland stem and progenitor cell activity after irradiation. Sci. Signal. 18, eady0398 (2025).

53. P. E. Burger, X. Xiong, S. Coetzee, S. N. Salm, D. Moscatelli, K. Goto, E. L. Wilson, Sca-1 expression identifies stem cells in the proximal region of prostatic ducts with high capacity to reconstitute prostatic tissue. Proc. Natl. Acad. Sci. 102, 7180–7185 (2005).

54. K. C. Bradley, K. Finsterbusch, D. Schnepf, S. Crotta, M. Llorian, S. Davidson, S. Y. Fuchs, P. Staeheli, A. Wack, Microbiota-Driven Tonic Interferon Signals in Lung Stromal Cells Protect from Influenza Virus Infection. Cell Rep. 28, 245-256.e4 (2019).

55. Q. Chen, H. Hirai, M. Chan, J. Zhang, M. Cho, S. H. Randell, P. K. L. Murthy, J. Rehman, Y. Liu, Characterization of perivascular alveolar epithelial stem cells and their niche in lung homeostasis and cancer. Stem Cell Rep. 19, 890–905 (2024).

56. X. Wu, V. L. D. Thi, Y. Huang, E. Billerbeck, D. Saha, H.-H. Hoffmann, Y. Wang, L. A. V. Silva, S. Sarbanes, T. Sun, L. Andrus, Y. Yu, C. Quirk, M. Li, M. R. MacDonald, W. M. Schneider, X. An, B. R. Rosenberg, C. M. Rice, Intrinsic Immunity Shapes Viral Resistance of Stem Cells. Cell 172, 423-438.e25 (2018).

57. J. Li, S. M. Dang, S. Sengupta, P. Schurmann, A. F. M. Dost, A. L. Moye, M. F. Trovero, S. Ahmed, M. Paschini, P. J. Bhetariya, R. Bronson, S. J. H. Sui, C. F. Kim, Organoid modeling reveals the tumorigenic potential of the alveolar progenitor cell state. EMBO J. 44, 1804–1828 (2025).

58. F. S. Rodrigues, A. Karoutas, S. Ruhland, N. Rabas, T. Rizou, S. D. Blasio, R. M. M. Ferreira, V. L. Bridgeman, R. Goldstone, M. L. Sopena, J.-H. Lee, L. Ombrato, I. Malanchi, Bidirectional activation of stem-like programs between metastatic cancer and alveolar type 2 cells within the niche. Dev. Cell, doi: 10.1016/j.devcel.2024.05.020 (2024).

59. R. Dagher, A. M. Copenhaver, V. Besnard, A. Berlin, F. Hamidi, M. Maret, J. Wang, X. Qu, Y. Shrestha, J. Wu, G. Gautier, R. Raja, M. Aubier, R. Kolbeck, A. A. Humbles, M. Pretolani, IL-33-ST2 axis regulates myeloid cell differentiation and activation enabling effective club cell regeneration. Nature Communications 11 (2020).

60. A. V. Misharin, L. Morales-Nebreda, P. A. Reyfman, C. M. Cuda, J. M. Walter, A. C. McQuattie-Pimentel, C.-I. Chen, K. R. Anekalla, N. Joshi, K. J. N. Williams, H. Abdala-Valencia, T. J. Yacoub, M. Chi, S. Chiu, F. J. Gonzalez-Gonzalez, K. Gates, A. P. Lam, T. T. Nicholson, P. J. Homan, S. Soberanes, S. Dominguez, V. K. Morgan, R. Saber, A. Shaffer, M. Hinchcliff, S. A. Marshall, A. Bharat, S. Berdnikovs, S. M. Bhorade, E. T. Bartom, R. I. Morimoto, W. E. Balch, J. I. Sznajder, N. S. Chandel, G. M. Mutlu, M. Jain, C. J. Gottardi, B. D. Singer, K. M. Ridge, N. Bagheri, A. Shilatifard, G. R. S. Budinger, H. Perlman, Monocyte-derived alveolar macrophages drive lung fibrosis and persist in the lung over the life span. Journal of Experimental Medicine 214, 2387–2404 (2017).

61. N. Joshi, S. Watanabe, R. Verma, R. P. Jablonski, C. I. Chen, P. Cheresh, N. S. Markov, P. A. Reyfman, A. C. McQuattie-Pimentel, L. Sichizya, Z. Lu, R. Piseaux-Aillon, D. Kirchenbuechler, A. S. Flozak, C. J. Gottardi, C. M. Cuda, H. Perlman, M. Jain, D. W. Kamp, G. R. S. Budinger, A. V. Misharin, A spatially restricted fibrotic niche in pulmonary fibrosis is sustained by M-CSF/M-CSFR signalling in monocyte-derived alveolar macrophages. European Respiratory Journal 55 (2020).

62. D. Aran, A. P. Looney, L. Liu, E. Wu, V. Fong, A. Hsu, S. Chak, R. P. Naikawadi, P. J. Wolters, A. R. Abate, A. J. Butte, M. Bhattacharya, Reference-based analysis of lung single-cell sequencing reveals a transitional profibrotic macrophage. Nature Immunology 20, 163–172 (2019).

63. T. Satoh, K. Nakagawa, F. Sugihara, R. Kuwahara, M. Ashihara, F. Yamane, Y. Minowa, K. Fukushima, I. Ebina, Y. Yoshioka, A. Kumanogoh, S. Akira, Identification of an atypical monocyte and committed progenitor involved in fibrosis. Nature 541, 96–101 (2017).

64. S. A. Eming, T. A. Wynn, P. Martin, Inflammation and metabolism in tissue repair and regeneration. Science 356, 1026–1030 (2017).

65. W. T’Jonck, C. C. Bain, The role of monocyte-derived macrophages in the lung: It’s all about context. Int. J. Biochem. Cell Biol. 159, 106421 (2023).

66. N. Zhang, R. S. Czepielewski, N. N. Jarjour, E. C. Erlich, E. Esaulova, B. T. Saunders, S. P. Grover, A. C. Cleuren, G. J. Broze, B. T. Edelson, N. Mackman, B. H. Zinselmeyer, G. J. Randolph, Expression of factor V by resident macrophages boosts host defense in the peritoneal cavity. J. Exp. Med. 216, 1291–1300 (2019).

67. A. S. Neupane, M. Willson, A. K. Chojnacki, F. V. E. S. Castanheira, C. Morehouse, A. Carestia, A. E. Keller, M. Peiseler, A. DiGiandomenico, M. M. Kelly, M. Amrein, C. Jenne, A. Thanabalasuriar, P. Kubes, Patrolling Alveolar Macrophages Conceal Bacteria from the Immune System to Maintain Homeostasis. Cell 183, 110-125.e11 (2020).

68. X. Zhang, S. Li, I. Malik, M. H. Do, L. Ji, C. Chou, W. Shi, K. J. Capistrano, J. Zhang, T.-W. Hsu, B. G. Nixon, K. Xu, X. Wang, A. Ballabio, L. S. Schmidt, W. M. Linehan, M. O. Li, Reprogramming tumour-associated macrophages to outcompete cancer cells. Nature 619, 616–623 (2023).

69. N. Almanzar, D. Yang, J. Xia, S. Udit, P. Joshi, S. Adhikari, D. A. Hoagland, S. T. Yeung, C. Khairallah, T. Huerta, A. Wallrapp, B. D. Umans, N. Sarden, O. Erdogan, N. Baalbaki, J. Hou, A. Beekmayer-Dhillon, J. Lee, K. A. Meerschaert, S. D. Liberles, R. A. Franklin, B. G. Yipp, K. M. Khanna, P. Baral, A. L. Haber, I. M. Chiu, Vagal TRPV1+ sensory neurons protect against influenza virus infection by regulating lung myeloid cell dynamics. Sci. Immunol. 10, eads6243 (2025).

70. D. A. Hoagland, R. A. Franklin, Isolation of Live Myeloid and Epithelial Cell Populations from the Mouse Lung. J. Vis. Exp. : JoVE, 10.3791/67648 (2025).

