## Supplementary Materials for "Type I interferon primes the alveolar epithelium to receive reparative signals from tissue-resident macrophages"

### Supplementary information

This file includes:

Figures S1 to S7
Tables S1 to S3

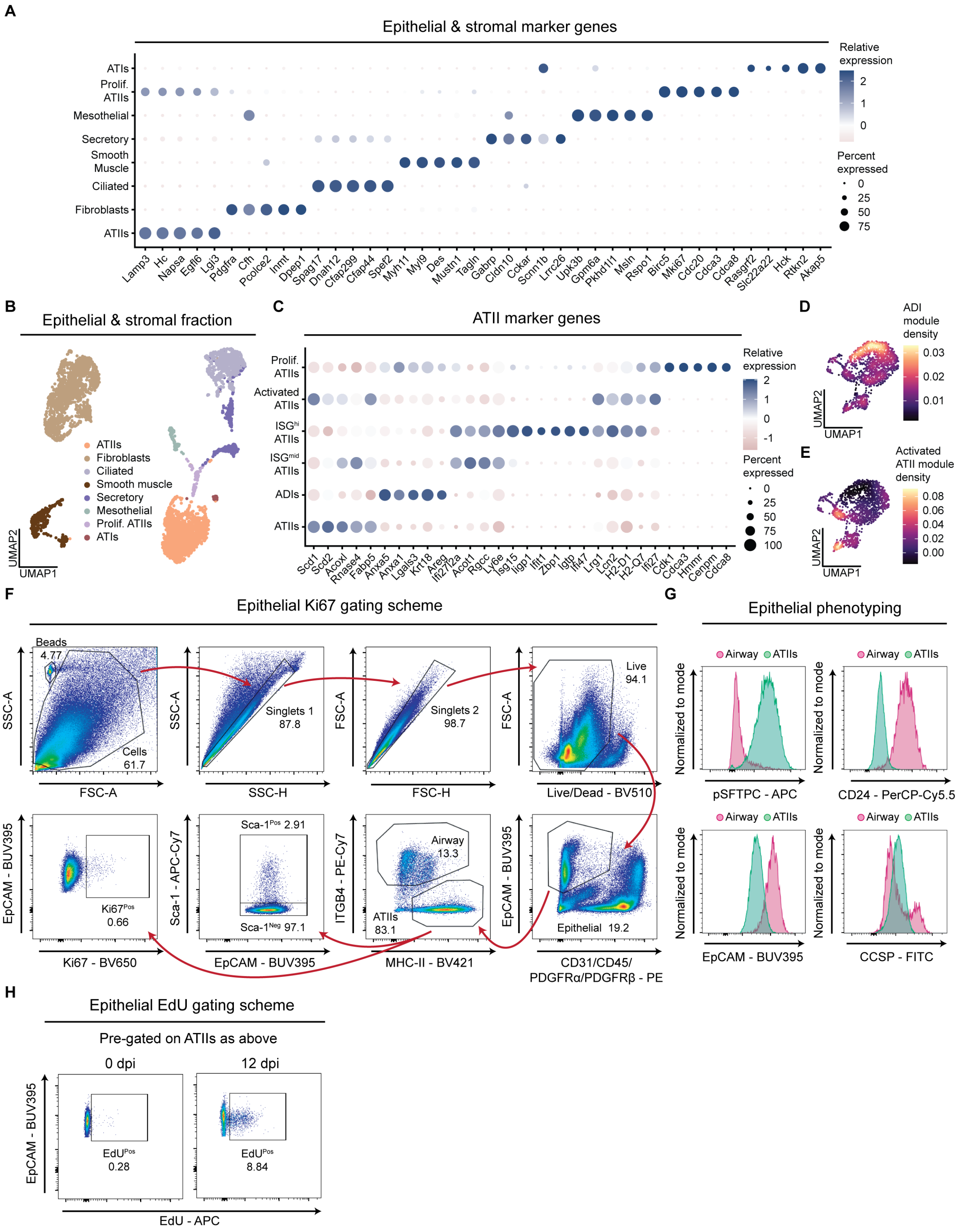


**Figure S1. Phenotypic profiling defines ISG$^{hi}$ Sca-1$^{Pos}$ ATIIs as a distinct lung epithelial population.** (**A–C**) Single-cell RNA-seq characterization of lung stromal and epithelial cells during sublethal i.n. IAV infection (A/PR/8/34; 6.5 TCID$_{50}$ for females and 32 TCID$_{50}$ for males). Mice were mock- (0 dpi) or IAV-infected and harvested at 7, 12, or 28 dpi. Dot plot of marker genes for epithelial and stromal populations (A; n = 1 male and 1 female mouse per group). UMAP of epithelial and stromal populations (4,907 cells; B). Dot plot of marker genes for ATII subclusters (C). (**D–E**) Nebulosa density plots of ADI (D) and activated ATII gene modules (E). (**F–H**) Flow cytometry gating and phenotyping of ATIIs (mock-infected). Gating scheme (F). Representative histograms of pSFTPC, CD24, EpCAM, and CCSP on ATIIs and airway cells as gated in F (G). Representative EdU gating scheme at 0 and 9 dpi (H).

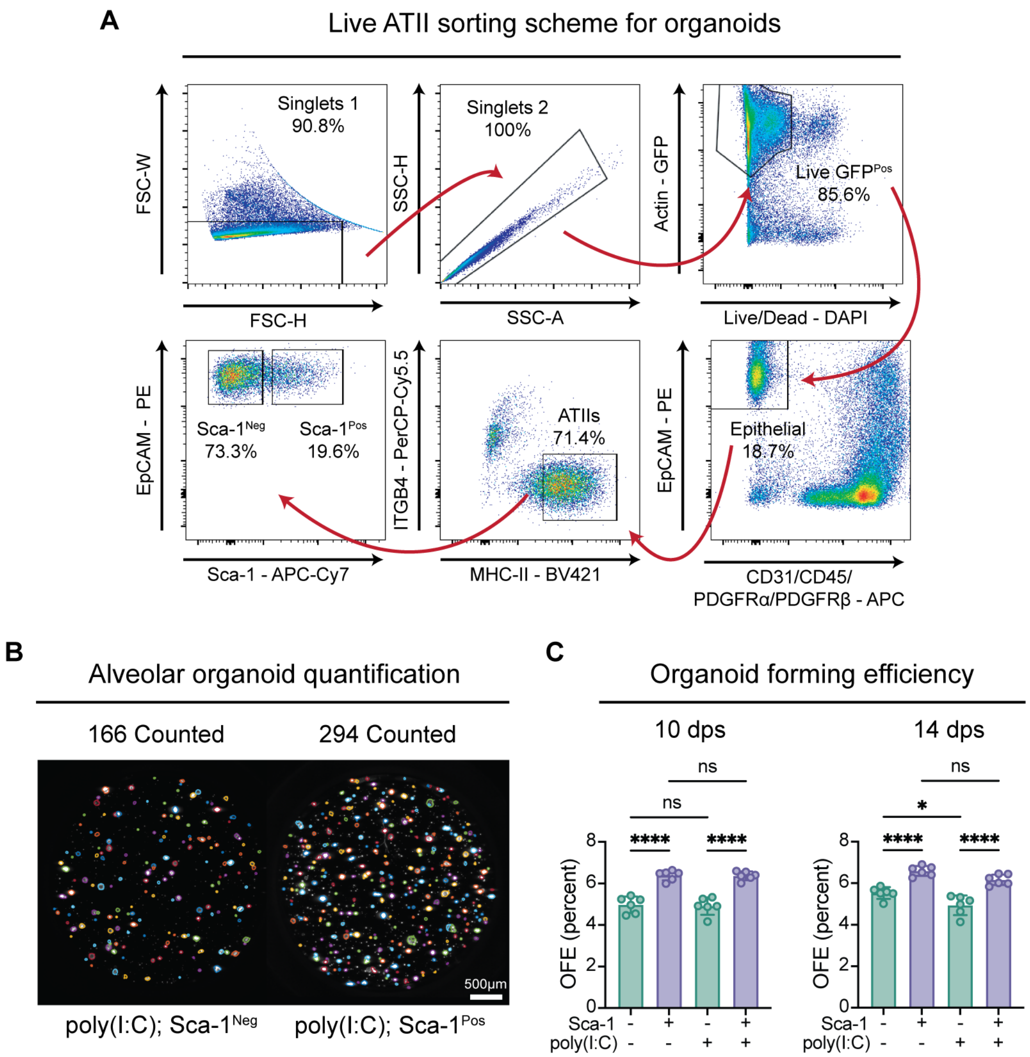


**Figure S2. Sca-1$^{Pos}$ ATIIs display enhanced progenitor capacity during *ex vivo* organoid culture. (A)** Representative gating scheme for live Sca-1$^{Neg/Pos}$ ATII sorting for organoid seeding from a PBS treated sample. (**B**) Representative MATLAB-processed images used for organoid quantification (7 dps) following poly(I:C) challenge (33.75 µg). Individual organoids counted are circled in varying colors for representation purposes. (**C**) Organoid-forming efficiency (OFE) of sorted PBS or poly(I:C) (33.75 µg)-challenged Sca-1$^{Neg/Pos}$ ATII-derived organoids at 10 and 14 dps (n = 2 biological replicates and 3 technical replicates per group). Data are representative of at least two independent experiments. One-way ANOVA with Dunnet's post-test for (C). Error bars indicate standard deviation (SD). $*p \leq 0.05, ****p \leq 0.0001$.

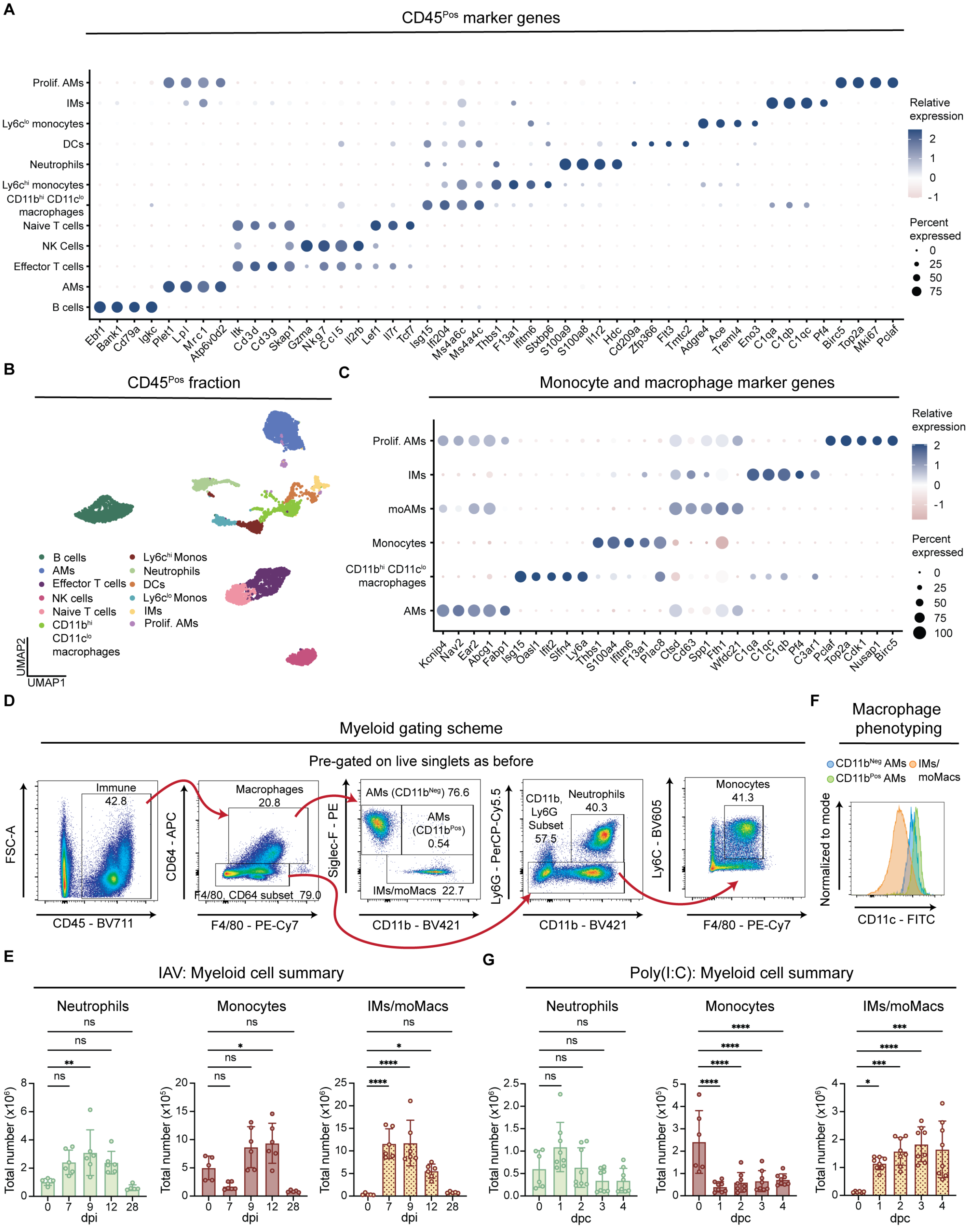


**Figure S3. Phenotypic profiling defines CD11b$^{Pos}$ AMs as a distinct lung myeloid population.** (**A–C**) CITE-seq characterization of immune cells during sublethal i.n. IAV infection (A/PR/8/34; 6.5 $TCID_{50}$ for females and 32 $TCID_{50}$ for males). Mice were mock- (0 dpi) or IAV-infected and harvested at 7, 12, or 28 dpi. Dot plot of marker genes for immune populations (A; n = 1 male and 1 female mouse per group). UMAP visualization of immune populations (7,300 cells; B). Dot plot of marker genes for macrophage and monocyte subclusters (C). (**D**) Flow cytometry gating of myeloid cells (mock-infected). (**E**) Quantification of neutrophils, monocytes, and IMs/moMacs during IAV infection (5 $TCID_{50}$; n = 5 - 6 mice per group). (**F**) Representative histogram of CD11c expression on macrophages (9 dpi IAV; 5 $TCID_{50}$). (**G**) Quantification of neutrophils, monocytes, and IMs/moMacs during poly(I:C) challenge (33.75 µg; n = 6 - 8 mice per group). Data are representative of at least two independent experiments. One-way ANOVA for (E), and (G). $*p \leq 0.05, **p \leq 0.005, ***p \leq 0.0005, ****p \leq 0.0001$.

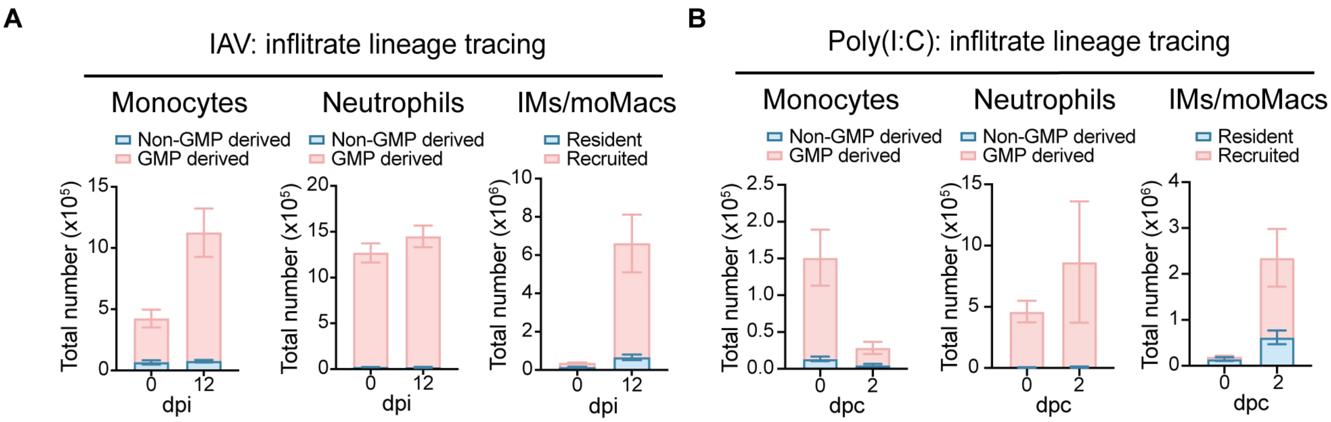


**Figure S4. Ms4a3$^{Cre}$ LSL-tdTomato mice label myeloid infiltrates of GMP origin across viral challenges.** (**A**) Flow cytometry quantification of monocytes, neutrophils and IM/moMacs 12 d following sublethal i.n. IAV infection (A/PR/8/34; 6.5 $TCID_{50}$; n = 3 - 4 mice per group). (**B**) Flow cytometry quantification of monocytes, neutrophils and IM/moMacs 2 d following i.t. poly(I:C) challenge (33.75 µg; n = 3 mice per group). Resident tdTomato$^{Neg}$ (non-GMP derived) in blue and recruited (GMP derived) tdTomato$^{Pos}$ in red. Data are representative of at least two independent experiments.

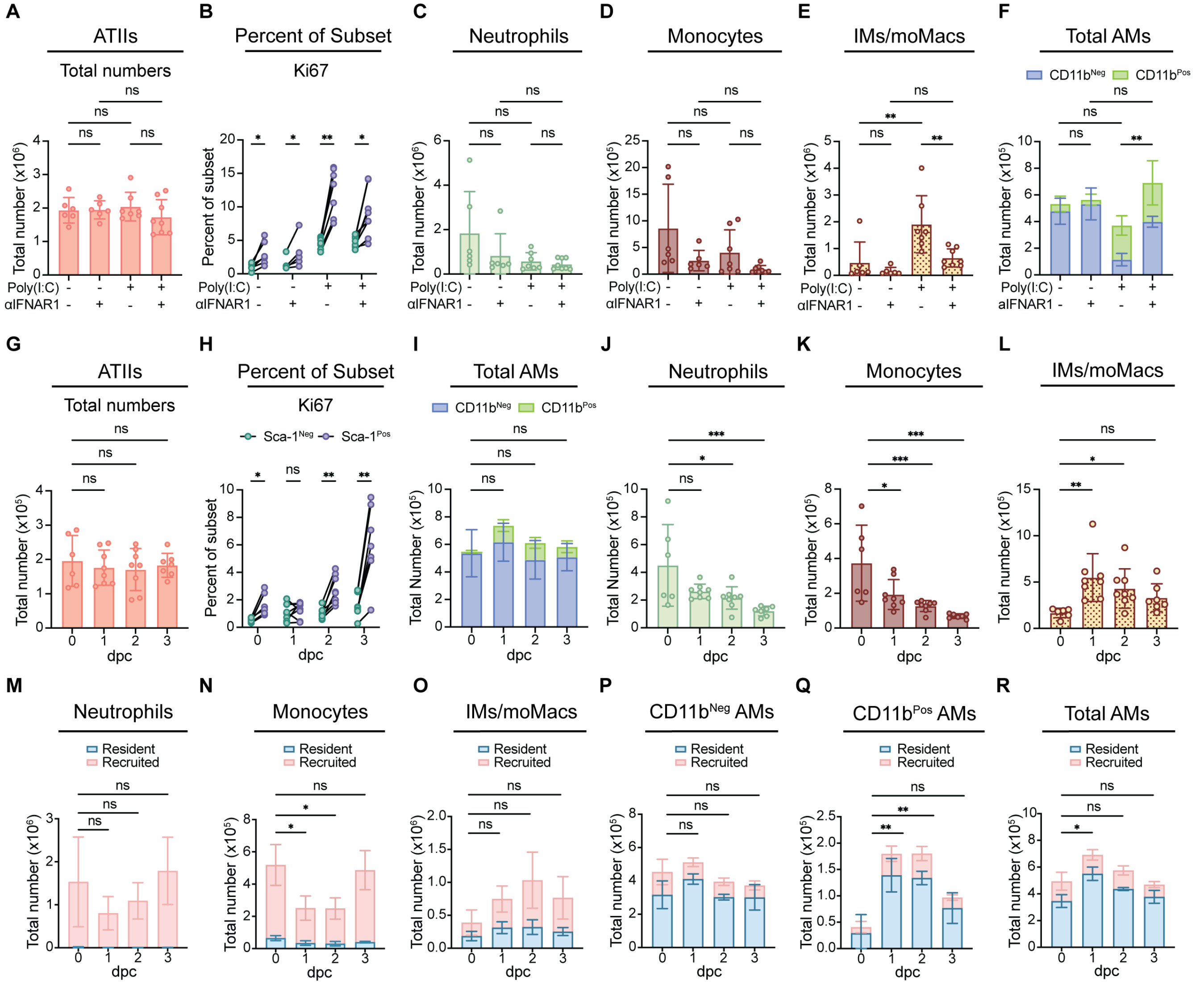


**Figure S5. IFN-I signaling alters lung myeloid cell compartment.** (**A–F**) Flow cytometry quantification of total ATIIs (A), percent Ki67$^{Pos}$ of Sca-1$^{Neg/Pos}$ ATIIs (B), neutrophils (C), monocytes (D), IMs/moMacs (E), and CD11b$^{Neg/Pos}$ AMs (F) following IFNAR1 blockade (i.t. 1x daily for 3 days; days 0, 1, 2; 200 µg per treatment) and harvested at 3 dpc (treated on day 0; i.t. 33.75 µg poly(I:C); n = 6 – 8 mice per group). (**G–L**) Flow cytometry quantification of total ATIIs (G) percent Ki67$^{Pos}$ of Sca-1$^{Neg/Pos}$ ATIIs (H), CD11b$^{Neg/Pos}$ AMs (I), neutrophils (J), monocytes (K), and IMs/moMacs (L) following exogenous IFN-I challenge (i.t. 1x daily for 2 days; 450 ng IFN-β per treatment; n = 6 – 8 mice per group). (**M–R**) Flow cytometry quantification of neutrophils (M), monocytes (N), IMs/moMacs (O), CD11b$^{Neg}$ AMs (P), CD11b$^{Pos}$ AMs (Q), and total AMs (R) after exogenous IFN-I challenge (i.t. 1x daily for 2 days; 700 ng IFN-β per treatment) in 12-16-wk-old *Ms4a3*$^{Cre}$ tdTomato mice (n = 3 mice per group). Resident tdTomato$^{Neg}$ in blue and recruited tdTomato$^{Pos}$ in red. Data from two independent experiments were pooled for (A–L). Data are representative of two independent experiments for (M–R). One-way ANOVA with Dunnet's post-test for (A–R), calculated on the sum of CD11b$^{Neg}$ and CD11b$^{Pos}$ AMs for (F) and (I), calculated on the sum of tdTomato$^{Neg}$ and tdTomato$^{Pos}$ populations for (M–R). $*p \leq 0.05, **p \leq 0.005, ***p \leq 0.0005$.

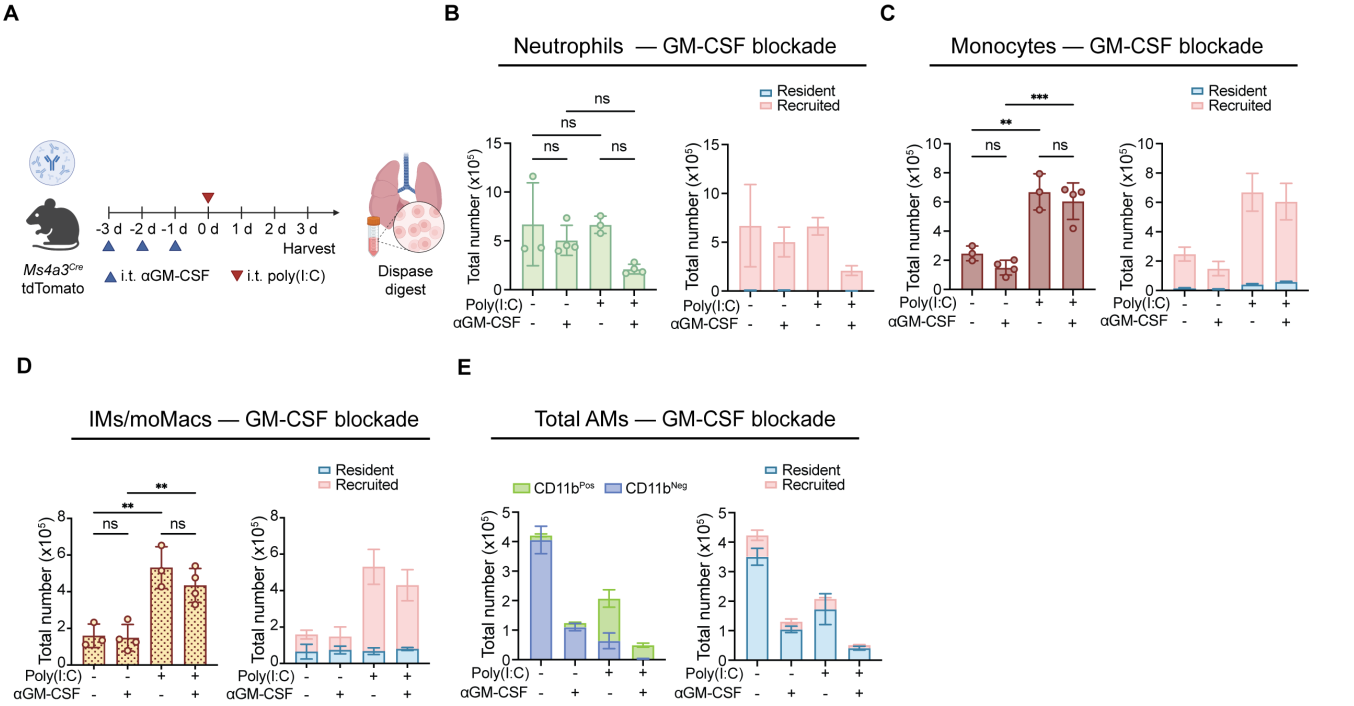


**Figure S6. GM-CSF blockade does not significantly deplete granulocyte monocyte progenitor (GMP)-derived myeloid infiltrates. (A)** Schematic of GM-CSF blockade (i.t. 1x daily for 3 days prior to challenge; 90 µg per treatment) in *Ms4a3*$^{Cre}$tdTomato mice at 3 d following poly(I:C) challenge (33.75 µg; n = 3 – 4 mice per group). **(B–E)** Flow cytometry quantification of neutrophils (B), monocytes (C), IMs/moMacs (D), and total AMs (E) and lineage depiction of each population. Resident tdTomato$^{Neg}$ in blue and recruited tdTomato$^{Pos}$ in red. One-way ANOVA with Dunnet's post-test for (B-D). $**p \leq 0.005, ***p \leq 0.0005$.

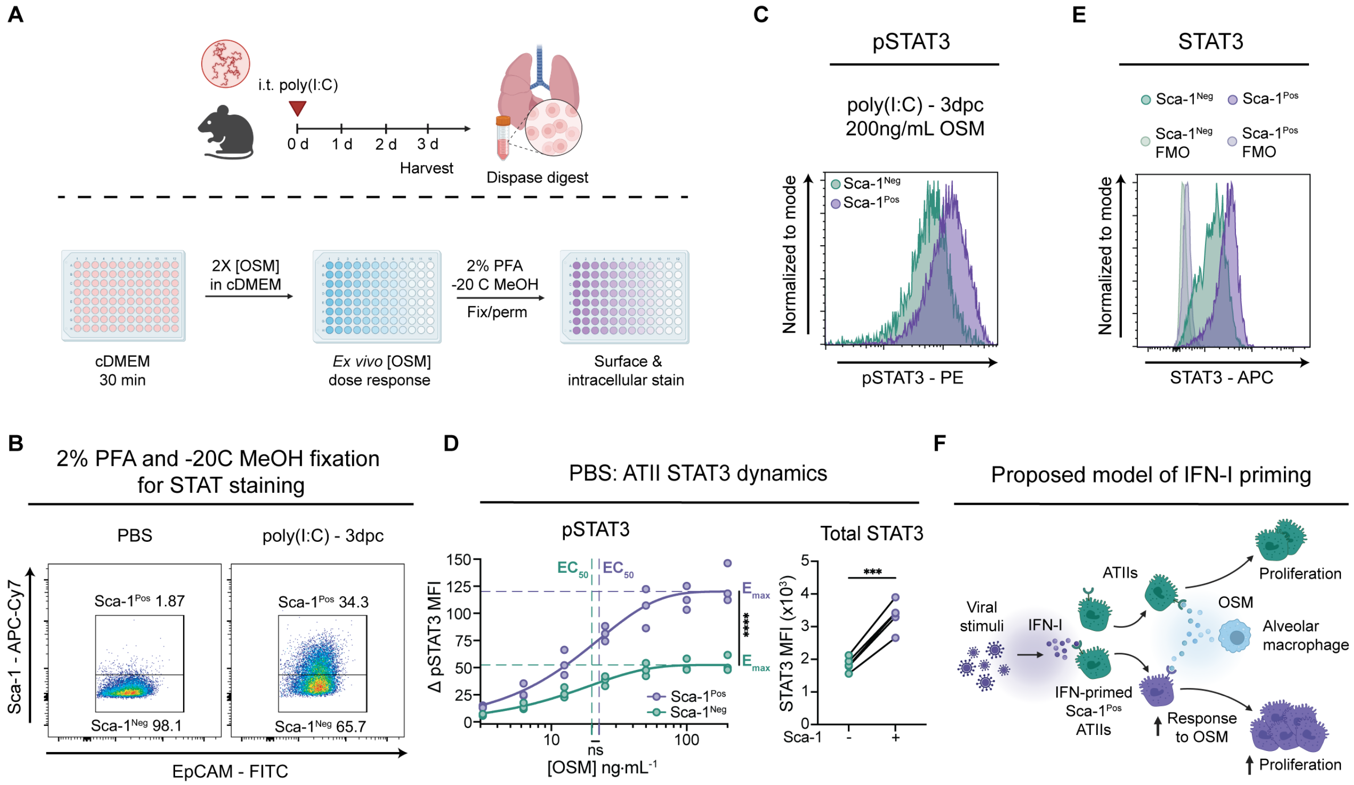


**Figure S7. IFN-primed Sca-1$^{Pos}$ ATIIs exhibit enhanced levels of pSTAT3 and STAT3.** **(A)** Poly(I:C) treatment scheme (33.75 µg) (top) and *ex vivo* OSM stimulation scheme of digested lung fractions (bottom). **(B)** Representative gating scheme for Sca-1$^{Neg/Pos}$ ATIIs from STAT3/pSTAT3 samples (2% PFA and -20 C MeOH fixed samples). **(C)** Representative histogram of pSTAT3. (**D**) Flow cytometry quantification of Δ phospho-STAT3 mean fluorescence intensity (pSTAT3 MFI) after mock challenge (0 dpc; PBS) followed by *ex vivo* OSM stimulation of digested lung fractions across a range of doses (mock, 3.125, 6.25, 12.5, 25, 50, 100, and 200 ng•mL$^{-1}$ with n = 3 biological replicates per OSM dose; calculated by subtracting out the average MFI of mock controls from the MFI of OSM stimulated samples) and total STAT3 MFI during mock challenge (0 dpc; PBS; n = 5 mice per group; calculated by subtracting out fluorescence minus one controls from raw MFI). (**E**) Representative histogram of STAT3. (**F**) Proposed model of IFN-I priming in ATIIs. Data are representative of at least two independent experiments. A one-phase association model was used to draw lines of best-fit for the pSTAT3 dose response curves and thereby determined $EC_{50}$ (tau) and $E_{max}$ (plateau) values. To test if $EC_{50}$ and $E_{max}$ values were different, sum-of-squares F tests were performed on these values. $***p \leq 0.0005, ****p \leq 0.0001$.

**Table S1. Top ten marker genes for each population in epithelial and stromal cell scRNA-seq and immune cell CITE-seq datasets.**

| Cluster | Gene | Mean Log2 FC | Adjusted p-values |
|---|---|---|---|
| ATIIs | *Lamp3* | 5.20 | 0 |
| | *Hc* | 5.20 | 0 |
| | *Napsa* | 5.19 | 0 |
| | *Egfl6* | 4.98 | 0 |
| | *Lgi3* | 4.95 | 0 |
| | *Etv5* | 4.70 | 0 |
| | *Ager* | 1.65 | 0 |
| | *S100g* | 4.94 | 0 |
| | *Cldn18* | 3.16 | 0 |
| | *Sfta2* | 4.70 | 0 |
| Fibroblasts | *Pdgfra* | 6.81 | 0 |
| | *Cfh* | 4.29 | 0 |
| | *Pcolce2* | 5.15 | 0 |
| | *Inmt* | 7.11 | 0 |
| | *Dpep1* | 7.49 | 0 |
| | *Ogn* | 3.95 | 0 |
| | *Pid1* | 6.73 | 0 |
| | *Cdh11* | 4.77 | 0 |
| | *Fn1* | 4.81 | 0 |
| | *Fbn1* | 4.00 | 0 |
| Ciliated | *Spag17* | 6.13 | 0 |
| | *Dnah12* | 6.31 | 0 |
| | *Cfap299* | 6.13 | 0 |
| | *Cfap44* | 6.08 | 0 |
| | *Spef2* | 6.14 | 0 |
| | *Cfap54* | 6.29 | 0 |
| | *Lrriq1* | 6.23 | 0 |
| | *Dnah5* | 6.25 | 0 |
| | *Cfap43* | 5.99 | 0 |
| | *Rp1* | 6.11 | 0 |
| Smooth muscle | *Myh11* | 9.02 | 0 |
| | *Myl9* | 6.53 | 0 |
| | *Des* | 5.85 | 0 |
| | *Mustn1* | 6.29 | 0 |
| | *Tagln* | 6.58 | 0 |
| | *Acta2* | 7.43 | 0 |
| | *Lmod1* | 7.31 | 0 |
| | *Tpm2* | 6.10 | 0 |
| | *Actg2* | 8.55 | 0 |
| | *Pde3a* | 5.11 | 0 |
| Secretory | *Gabrp* | 6.72 | 0 |
| | *Cldn10* | 4.62 | 0 |
| | *Cckar* | 3.77 | 0 |
| | *Scnn1b* | 3.31 | 0 |
| | *Lrrc26* | 6.61 | 0 |
| | *Mgat3* | 3.23 | 0 |
| | *Itprid1* | 6.89 | 0 |
| | *Fmo3* | 5.09 | 0 |
| | *Chad* | 6.45 | 0 |
| | *Tnfrsf21* | 3.17 | 0 |
| Mesothelial | *Upk3b* | 9.97 | 0 |
| | *Gpm6a* | 9.47 | 0 |
| | *Pkhd1l1* | 10.31 | 0 |
| | *Msln* | 7.86 | 0 |
| | *Rspo1* | 6.43 | 0 |
| | *Cldn15* | 8.61 | 0 |
| | *Wt1* | 7.38 | 0 |
| | *Aldh1a2* | 5.63 | 0 |
| | *Lrrn4* | 10.35 | 0 |
| | *Muc16* | 9.37 | 0 |
| Prolif. ATIIs | *Birc5* | 7.90 | 0 |
| | *Mki67* | 8.03 | 0 |
| | *Cdc20* | 6.88 | 0 |
| | *Cdca3* | 7.60 | 0 |
| | *Cdca8* | 7.45 | 0 |
| | *Cdk1* | 6.28 | 0 |
| | *Tpx2* | 6.84 | 0 |
| | *Cdkn3* | 7.89 | 0 |
| | *Cenpm* | 6.81 | 0 |
| | *Racgap1* | 7.02 | 0 |
| ATIs | *Rasgrf2* | 9.70 | 0 |
| | *Slc22a22* | 10.42 | 1.47E-244 |
| | *Hck* | 7.38 | 4.55E-221 |
| | *Rtkn2* | 9.28 | 1.01E-212 |
| | *Akap5* | 7.60 | 3.37E-143 |
| | *Ptpre* | 6.90 | 1.46E-112 |
| | *Scel* | 7.10 | 5.12E-112 |
| | *Sema3a* | 6.45 | 8.71E-78 |
| | *Cttnbp2* | 6.41 | 2.19E-73 |
| | *Flrt3* | 6.99 | 3.30E-73 |

| Cluster | Gene | Mean Log2 FC | Adjusted p-values |
|---|---|---|---|
| B cells | *Ebf1* | 6.74 | 0 |
| | *Bank1* | 6.34 | 0 |
| | *Cd79a* | 6.48 | 0 |
| | *Igkc* | 5.83 | 0 |
| | *Ighm* | 4.76 | 0 |
| | *Mef2c* | 4.48 | 0 |
| | *H2-Eb1* | 1.98 | 0 |
| | *Ly6d* | 6.53 | 0 |
| | *H2-Aa* | 2.06 | 0 |
| | *Aff3* | 3.14 | 0 |
| AMs | *Plet1* | 5.84 | 0 |
| | *Lpl* | 5.31 | 0 |
| | *Mrc1* | 4.46 | 0 |
| | *Atp6v0d2* | 5.32 | 0 |
| | *Ear2* | 4.56 | 0 |
| | *Kcnip4* | 6.06 | 0 |
| | *Tcf7l2* | 3.80 | 0 |
| | *Dst* | 4.58 | 0 |
| | *Chil3* | 4.88 | 0 |
| | *Ltc4s* | 5.23 | 0 |
| Effector T cells | *Itk* | 2.69 | 0 |
| | *Cd3d* | 3.37 | 0 |
| | *Cd3g* | 3.86 | 0 |
| | *Skap1* | 2.43 | 0 |
| | *Trbc2* | 2.86 | 0 |
| | *Ms4a4b* | 2.53 | 0 |
| | *Ptpn22* | 2.57 | 0 |
| | *Themis* | 3.38 | 0 |
| | *Trac* | 3.60 | 0 |
| | *Icos* | 4.32 | 0 |
| NK cells | *Gzma* | 6.36 | 0 |
| | *Nkg7* | 3.75 | 0 |
| | *Ccl5* | 3.32 | 0 |
| | *Il2rb* | 4.19 | 0 |
| | *Il12rb2* | 4.47 | 0 |
| | *Prf1* | 6.09 | 0 |
| | *AW112010* | 3.20 | 0 |
| | *Klre1* | 6.30 | 0 |
| | *Arsb* | 4.36 | 0 |
| | *Klrd1* | 3.50 | 0 |
| Naive T cells | *Lef1* | 3.81 | 0 |
| | *Itk* | 2.44 | 0 |
| | *Il7r* | 3.41 | 0 |
| | *Tcf7* | 4.37 | 0 |
| | *Camk4* | 2.99 | 0 |
| | *Dapl1* | 6.18 | 1.53E-295 |
| | *Dusp10* | 3.85 | 3.36E-290 |
| | *Trbc2* | 2.62 | 5.44E-277 |
| | *Gm2682* | 2.35 | 3.57E-272 |
| | *Rps29* | 1.26 | 3.06E-269 |
| CD11b$^{hi}$ CD11c$^{lo}$ AMs | *Isg15* | 3.58 | 0 |
| | *Ifi204* | 3.63 | 0 |
| | *Ms4a6c* | 3.27 | 0 |
| | *Ms4a4c* | 3.52 | 0 |
| | *Irf7* | 3.28 | 0 |
| | *Ccl2* | 4.87 | 0 |
| | *Gbp2* | 3.08 | 0 |
| | *Mafb* | 3.69 | 0 |
| | *Oasl2* | 3.24 | 0 |
| | *Fcgr1* | 4.39 | 0 |
| Ly6C$^{hi}$ Monocytes | *Thbs1* | 3.79 | 0 |
| | *F13a1* | 5.42 | 0 |
| | *Ifitm6* | 3.88 | 0 |
| | *Stxbp6* | 4.76 | 0 |
| | *S100a4* | 2.83 | 5.20E-267 |
| | *Ccr2* | 3.15 | 2.00E-247 |
| | *Ifitm3* | 2.57 | 1.20E-241 |
| | *Ccl9* | 3.62 | 1.90E-235 |
| | *Emilin2* | 2.84 | 5.72E-228 |
| | *Plcb1* | 2.94 | 1.65E-206 |
| Neutrophils | *S100a9* | 8.15 | 0 |
| | *S100a8* | 8.15 | 0 |
| | *Il1r2* | 6.39 | 0 |
| | *Hdc* | 7.02 | 0 |
| | *Il1b* | 5.52 | 0 |
| | *Retnlg* | 8.52 | 0 |
| | *Csf3r* | 5.96 | 0 |
| | *Slc7a11* | 5.67 | 0 |
| | *Clec4d* | 5.36 | 0 |
| | *G0s2* | 7.23 | 0 |

| Cluster | Gene | Mean Log2 FC | Adjusted p-values |
|---|---|---|---|
| DCs | *Cd209a* | 6.76 | 0 |
| | *Zfp366* | 6.66 | 0 |
| | *Flt3* | 4.64 | 3.36E-267 |
| | *Tmtc2* | 5.03 | 1.21E-266 |
| | *Ctnnd2* | 6.18 | 3.91E-265 |
| | *Il4i1* | 5.38 | 3.61E-247 |
| | *Tmem176a* | 3.81 | 4.95E-226 |
| | *Htr7* | 4.51 | 1.07E-223 |
| | *Ccnd1* | 4.42 | 7.68E-203 |
| | *Tmem176b* | 3.47 | 5.55E-201 |
| Ly6c$^{lo}$ Monocytes | *Adgre4* | 6.30 | 0 |
| | *Ace* | 5.60 | 0 |
| | *Treml4* | 4.82 | 2.23E-254 |
| | *Eno3* | 5.22 | 1.51E-235 |
| | *Cd300e* | 5.94 | 1.15E-206 |
| | *Rap1gap2* | 3.52 | 2.44E-185 |
| | *Tiam2* | 4.76 | 2.25E-163 |
| | *Pot1b* | 4.17 | 4.24E-162 |
| | *Ldlrad3* | 3.92 | 4.31E-151 |
| | *Ceacam1* | 4.55 | 6.99E-148 |
| IMs | *C1qa* | 5.88 | 0 |
| | *C1qb* | 5.28 | 0 |
| | *C1qc* | 5.78 | 0 |
| | *Pf4* | 7.00 | 0 |
| | *Ms4a7* | 4.65 | 1.18E-268 |
| | *C3ar1* | 4.03 | 9.04E-258 |
| | *Stab1* | 5.40 | 2.03E-208 |
| | *Cxcl16* | 2.80 | 1.21E-157 |
| | *Tmem176b* | 3.69 | 1.54E-155 |
| | *Ccl8* | 5.93 | 3.87E-153 |
| Prolif. AMs | *Birc5* | 7.29 | 0 |
| | *Top2a* | 6.40 | 0 |
| | *Mki67* | 5.87 | 0 |
| | *Pclaf* | 7.63 | 0 |
| | *Cdk1* | 7.25 | 0 |
| | *Nusap1* | 6.27 | 0 |
| | *Prc1* | 5.75 | 0 |
| | *Cenpe* | 5.98 | 0 |
| | *Ube2c* | 7.50 | 0 |
| | *Cdca3* | 6.34 | 0 |

**Table S2. Top ten marker genes for each population in ATII scRNA-seq, and macrophage CITE-seq subsets.**

| Cluster | Gene | Mean Log2 FC | Adjusted p-values |
|---|---|---|---|
| ATIIs | *Scd1* | 0.92 | 2.00E-53 |
| | *Scd2* | 0.88 | 2.24E-53 |
| | *Acoxl* | 1.08 | 4.35E-44 |
| | *Rnase4* | 0.52 | 5.01E-29 |
| | *Fabp5* | 0.66 | 2.51E-28 |
| | *Abca3* | 0.43 | 1.05E-27 |
| | *Lgi3* | 0.52 | 1.17E-27 |
| | *Hc* | 0.54 | 2.01E-25 |
| | *Cbr2* | 0.41 | 3.14E-25 |
| | *Acly* | 0.65 | 3.90E-23 |
| ADIs | *Anxa5* | 1.81 | 1.59E-55 |
| | *Anxa1* | 2.34 | 5.74E-47 |
| | *Lgals3* | 2.79 | 4.52E-46 |
| | *Krt18* | 1.59 | 3.93E-45 |
| | *Areg* | 2.40 | 5.34E-40 |
| | *Ndnf* | 2.74 | 2.81E-37 |
| | *S100a6* | 4.18 | 1.51E-36 |
| | *Anxa3* | 1.29 | 8.79E-34 |
| | *Thbs1* | 2.70 | 3.82E-31 |
| | *Krt8* | 1.50 | 8.91E-31 |
| ISG$^{hi}$ ATIIs | *Ifi27l2a* | 4.17 | 6.84E-131 |
| | *Isg15* | 3.82 | 3.03E-118 |
| | *Ly6e* | 1.67 | 2.17E-86 |
| | *Bst2* | 2.52 | 2.24E-75 |
| | *Rgcc* | 2.05 | 1.02E-73 |
| | *Rtp4* | 3.21 | 3.62E-72 |
| | *Acot1* | 2.17 | 4.14E-72 |
| | *Irf7* | 3.61 | 1.67E-69 |
| | *Oasl2* | 3.64 | 1.32E-67 |
| | *Xaf1* | 2.95 | 2.06E-67 |
| Activated ATIIs | *Lrg1* | 1.95 | 6.79E-37 |
| | *Lcn2* | 1.19 | 1.13E-36 |
| | *H2-D1* | 1.10 | 7.78E-33 |
| | *H2-Q7* | 1.65 | 7.19E-28 |
| | *Ifi27* | 1.29 | 5.45E-26 |
| | *Ptgs1* | 1.26 | 6.67E-26 |
| | *H2-K1* | 0.93 | 1.98E-25 |
| | *Gapdh* | 0.86 | 2.06E-25 |
| | *Atp5pb* | 0.69 | 3.77E-23 |
| | *Cxcl15* | 0.70 | 4.49E-23 |
| Prolif. ATIIs | *Cdk1* | 8.78 | 2.14E-213 |
| | *Cdca3* | 8.67 | 3.38E-203 |
| | *Hmmr* | 10.22 | 2.48E-192 |
| | *Cenpm* | 7.05 | 9.72E-191 |
| | *Cdca8* | 8.31 | 2.55E-189 |
| | *Birc5* | 7.07 | 3.97E-189 |
| | *Pbk* | 9.59 | 6.44E-183 |
| | *Top2a* | 9.05 | 6.63E-179 |
| | *Mki67* | 7.90 | 1.09E-173 |
| | *Ckap2* | 8.09 | 1.34E-169 |

| Cluster | Gene | Mean Log2 FC | Adjusted p-values |
|---|---|---|---|
| AMs | *Kcnip4* | 2.98 | 8.47E-264 |
| | *Nav2* | 3.16 | 2.11E-225 |
| | *Ear2* | 2.30 | 6.63E-208 |
| | *Abcg1* | 1.95 | 2.25E-198 |
| | *Fabp1* | 3.78 | 1.11E-191 |
| | *Plet1* | 1.85 | 6.94E-182 |
| | *Ear1* | 3.13 | 1.48E-176 |
| | *Lpl* | 1.74 | 2.72E-172 |
| | *Tcf7l2* | 1.69 | 9.91E-169 |
| | *Cd9* | 1.61 | 1.59E-168 |
| CD11b$^{hi}$ CD11c$^{lo}$ AMs | *Isg15* | 6.06 | 1.07E-262 |
| | *Oasl1* | 5.97 | 7.51E-213 |
| | *Ifit2* | 6.46 | 8.35E-211 |
| | *Slfn4* | 6.34 | 2.01E-210 |
| | *Ly6a* | 5.72 | 1.06E-208 |
| | *Iigp1* | 6.65 | 4.31E-206 |
| | *Irf7* | 4.71 | 5.28E-206 |
| | *Spon1* | 7.00 | 1.48E-197 |
| | *Ms4a4c* | 4.14 | 7.46E-190 |
| | *Ifi205* | 5.88 | 8.08E-184 |
| Monocytes | *Thbs1* | 4.84 | 2.45E-226 |
| | *S100a4* | 3.55 | 1.04E-208 |
| | *Ifitm6* | 4.54 | 1.10E-202 |
| | *F13a1* | 3.95 | 1.30E-182 |
| | *Plac8* | 2.21 | 5.91E-169 |
| | *Stxbp6* | 5.49 | 1.41E-153 |
| | *S100a6* | 2.32 | 7.28E-148 |
| | *Adgre5* | 3.40 | 1.88E-135 |
| | *Ifitm3* | 2.11 | 3.36E-134 |
| | *Ccr2* | 2.86 | 1.62E-133 |
| moAMs | *Ctsd* | 1.54 | 3.49E-75 |
| | *Cd63* | 1.78 | 5.17E-67 |
| | *Spp1* | 2.13 | 2.03E-63 |
| | *Fth1* | 1.17 | 3.25E-63 |
| | *Wfdc21* | 1.76 | 2.80E-55 |
| | *Ccl6* | 1.32 | 5.14E-54 |
| | *Fabp5* | 1.50 | 4.07E-51 |
| | *Ftl1* | 1.00 | 6.61E-47 |
| | *Acp5* | 1.66 | 6.81E-47 |
| | *Lyz2* | 1.09 | 1.21E-43 |
| IMs | *C1qa* | 4.94 | 2.62E-165 |
| | *C1qc* | 4.85 | 7.31E-162 |
| | *C1qb* | 4.25 | 7.15E-148 |
| | *Pf4* | 6.56 | 1.58E-128 |
| | *C3ar1* | 2.92 | 2.88E-74 |
| | *Ms4a7* | 3.48 | 1.94E-71 |
| | *Apoe* | 4.73 | 1.05E-70 |
| | *Tmem176b* | 3.53 | 4.85E-66 |
| | *Tmem176a* | 4.02 | 6.85E-61 |
| | *Blnk* | 4.29 | 4.49E-56 |

| Cluster | Gene | Mean Log2 FC | Adjusted p-values |
|---|---|---|---|
| Prolif. AMs | *Pclaf* | 7.54 | 1.47E-283 |
| | *Top2a* | 7.21 | 4.85E-278 |
| | *Cdk1* | 6.43 | 5.77E-244 |
| | *Nusap1* | 6.25 | 1.13E-239 |
| | *Birc5* | 6.58 | 1.18E-239 |
| | *Rrm2* | 7.59 | 1.64E-220 |
| | *Prc1* | 6.27 | 1.08E-216 |
| | *Cenpe* | 6.91 | 2.71E-213 |
| | *Kif15* | 6.54 | 2.59E-210 |
| | *Cdca8* | 6.22 | 2.42E-204 |

**Table S3. Bespoke gene modules adapted from the literature used for ATII subtype annotation.**

| Gene Module | Genes | Adapted from (PMID): |
|---|---|---|
| ADI module | *Krt8* , *Cdkn1a* , *Hbegf* , *Areg* , *Itgb6* | 32750316 32678092 |
| ISG module | *Isg15* , *Oasl2* , *Ifitm3* , *Irf7* , *Stat1* , *Usp18* , *Stat2* , *Rtp4* | 30224793 |
| Activated ATII module | *Ifi27l2a* , *Lcn2* , *Il33* , *Retnla* , *Cxcl17* , *Lrg1* , *Itga7* , *Ptges* , *Glrx* | 32750316 32678092 |
